# Developmental delay ensures global tissue size robustness upon local induction of apoptosis

**DOI:** 10.1101/2024.12.21.629882

**Authors:** Ralitza Staneva, Gabriel Sobczyk-Moran, Florence Levillayer, Alexis Villars, Anđela Davidović, Romain Levayer

## Abstract

The capacity of our tissues to cope with external and internal stress relies on the tight coupling between cell proliferation, cell growth and cell death. This coupling is assumed to be based on compensatory proliferation, where local mitogenic signals and mechanical inputs generated by dying cells promote neighbouring cell proliferation. However, compensatory proliferation was mostly studied in the context of massive death induction, irradiation, surgical tissue ablation or upon genetic perturbation of apoptosis execution. It remains thus unclear whether the same mechanism operates during physiological programmed cell death or upon mild induction of apoptosis, especially *in vivo*. Here, we use the *Drosophila* prospective wing (the larval wing disc), to study the impact of local induction of apoptosis on tissue size and proliferation pattern. We first confirmed that the wing could recover its final size and compensate for mild induction of apoptosis. However, using spatial statistics we found surprisingly that local induction of death is not associated with any local increase of proliferation, could it be upon clonal or compartment induction of apoptosis. Compensation is driven instead by a JNK dependant delay of growth and lengthening of the larval stage which is required to reach the final tissue target size. These results suggest that compensation is here driven by a global response rather that a local proliferation induction. Accordingly, while total tissue size is maintained despite local induction of apoptosis, this mechanism fails to correct the local reduction of cell number, hence modulating wing shape and proportion. Overall, this study opens novel perspectives on tissue size regulation and outlines the context-dependency of compensatory mechanisms.

## Introduction

The robustness of organ and tissue size is based on their capacity to adapt to local and global perturbations and to fine tune the core parameters regulating cell number and cell size. For instance, the high rate of turnover in the gut, corresponding to billions of cell dying and dividing on a daily basis^1^, imposes a very tight coupling between the proliferation of stem cell and the extrusion of cells at gut villi. Alternatively, the precise regulation of growth and the final size of developing organs is mediated by the fine-tuning of cell size/volume, cell proliferation, and programmed cell death^2,3^. While there has been a lot of effort dissecting the contribution of cell growth and cell proliferation to organ size regulation^4,5^, the coupling between cell death and growth remained poorly understood, especially in physiological contexts.

Apoptosis-induced compensatory proliferation is the mechanism promoting cell proliferation in vicinity of dying apoptotic cells^6,7^. Most of our molecular knowledge of compensatory proliferation was obtained in the *Drosophila* wing imaginal disc, an epithelial sac that undergoes massive proliferation during larval stage and gives rise to the adult wing and thorax. Compensatory proliferation in the wing disc was initially characterised in the context of *Drosophila* larvae irradiation^8,9^ which outlined the capacity of the wing to compensate for massive cell loss. Similar mechanisms were characterised upon genetic induction of apoptosis in a large tissue domain^10^, surgery mediated tissue ablation^11,12^, or upon inhibition of the executioner caspases and accumulation of so-called “undead cells”^13,14^. Compensation is mostly mediated by the secretion of mitogenic factors (which are tissue and context dependent^14^) from the dying cells, which then promote cell proliferation in the neighbouring cells^14–17^. Similar principles were outlined in the context of head regeneration in hydra^18^, regeneration of *Drosophila* midgut^19^, Xenopus tadpole tail^20^,Zebrafish tail^21^ or upon perturbation of apoptosis execution in mammalian skin^22^, suggesting that compensatory mechanisms are rather well conserved. However, most of these perturbative approaches rely on acute elimination of a large proportion of tissue and/or perturbation of the execution of apoptosis. As such, it remains unclear whether the same signalings could occur in the context of physiological cell death or mild induction of apoptosis. Moreover, most of these perturbations rely on ubiquitous induction of apoptosis or ablation of large domains over long time scales, preventing quantitative characterisation of the spatial distribution of compensation. Recently, we outlined the existence of local hotspots of apoptosis in the wing disc which correlated with a local reduction of net clonal growth and tissue size, suggesting that this local physiological increase of apoptosis is not compensated by a local increase of growth/proliferation^2^. More recently, the coupling between single cell death/extrusion and proliferation was outlined in MDCK cells, where single cell extrusion is associated with neighbouring cell deformation and the activation of YAP/TAZ and mitosis in a subset of neighbouring cells^23^. Yet, whether this local response can also be observed *in vivo* and whether this could more generally apply to any epithelial tissue remains an open question.

Here, we use the *Drosophila* wing disc to test more generally whether proliferation is indeed coupled with apoptosis, especially in the context of relatively mild induction of apoptosis (elimination of ∼5% of the cells). We first confirmed that the adult wing could indeed compensate such mild loss of cells. However, to our great surprise, using various statistical and spatial analyses, we could never a find a significant increase of cell proliferation in vicinity of dying cells, nor any distinctive spatial features in the distribution of cell proliferation either in the context of apoptosis induction in clones or in a full compartment. Alternatively, we found an apparent transient and global increase of cell proliferation rate after apoptosis induction, which was mostly explained by the slow-down of tissue growth progression during the apoptosis induction phase and the higher proliferation rate in smaller discs. This transient slow-down was driven by JNK pathway activation in the dying cells which slightly delay the pupariation timing, most likely through the relaxin hormone Dilp8. This global modulation of developmental timing was required to compensate for the loss of cells and ensure the robustness of global tissue size. However, this global response fails to correct for local variations of tissue size driven by local induction of apoptosis. Altogether, this works outlines that apoptosis is not systematically associated with local compensatory proliferation, and that other mechanisms based on the alteration of developmental timing ensure global tissue size robustness, while being poorly efficient for the correction of local perturbations. This opens novel perspectives for the understanding of organ size and shape robustness.

## Results

### Local apoptosis does not trigger a local increase of proliferation

To study the impact of local induction of apoptosis on tissue growth and size, we used a conditional induction of the pro-apoptotic gene *reaper* in clones covering 2-12%, median at 5%, of the surface of the wing imaginal disc (**Fig. 1A, Fig. S1A**). Induction of the pro-apoptotic gene reaper for 8 or 16h at 96 hours after egg laying (at a developmental window still permissive for regeneration^10^) was sufficient to eliminate a large proportion of clonal cells (**Fig. 1B**). This small, albeit significant, cell loss was fully compensated in adult wings since their size was indistinguishable from controls wings (**Fig. 1C**), demonstrating that mild apoptosis can still be compensated. We therefore looked for the mechanism that could explain the recovery of tissue size. Compensatory proliferation is supposed to promote local proliferation through the secretion of mitogenic factors^6,7^. We therefore performed a systematic analysis of the spatial distribution of mitotic cells relative to apoptotic cells. Surprisingly, the local density of Phospho Histone 3 positive cells (a mitotic marker) was not increased in vicinity of apoptotic clones after 8 or 16h hours of apoptosis induction (**Fig. 1D,E, Fig. S1B),** as outlined by the analysis of PH3 density in concentric bands, the comparison with distribution of randomised PH3 localisation (**Fig. S1 D,E**), or by comparing the distribution of closest neighbours between clones and PH3 cells after randomisation of their position (**Fig. S1C**). To confirm that this absence is not due to a longer delay required for proliferation induction, spatial analysis was also performed after a recovery time of 5 or 10h after the end of apoptosis. Since most of the clone have disappeared after this recovery, we use an alternative spatial statistics method to reveal potential biases in the spatial distribution of mitoses. The Ripley K- and G-function are used to reveal bias in the spatial distribution relative to a random/Poisson process and can be used to outline clustering or dispersive effects^24–26^. We reasoned that a local induction of mitosis near apoptotic clones should locally increase mitosis concentration and lead to a more clustered distribution. However, none of the conditions we used for apoptosis induction leads to a significant increase mitosis clustering relative to control clones (**Fig. S2**). Finally, we confirmed the absence of local induction of proliferation by analysing the spatial pattern of proliferation in the posterior wing disc compartment upon apoptosis induction in the anterior compartment for 8 hours, immediately after death induction, and after 16 or 24h of recovery at 18° (**Fig. 1F,G, Fig. S3**). None of these conditions was associated with a local increase of proliferation and/or a spatial bias in mitotic cell localisation compared to control (**Fig. 1G, Fig. S3**). Altogether, these results suggest that mild induction of apoptosis does not lead to a significant local increase of proliferation.

**Figure 1:**
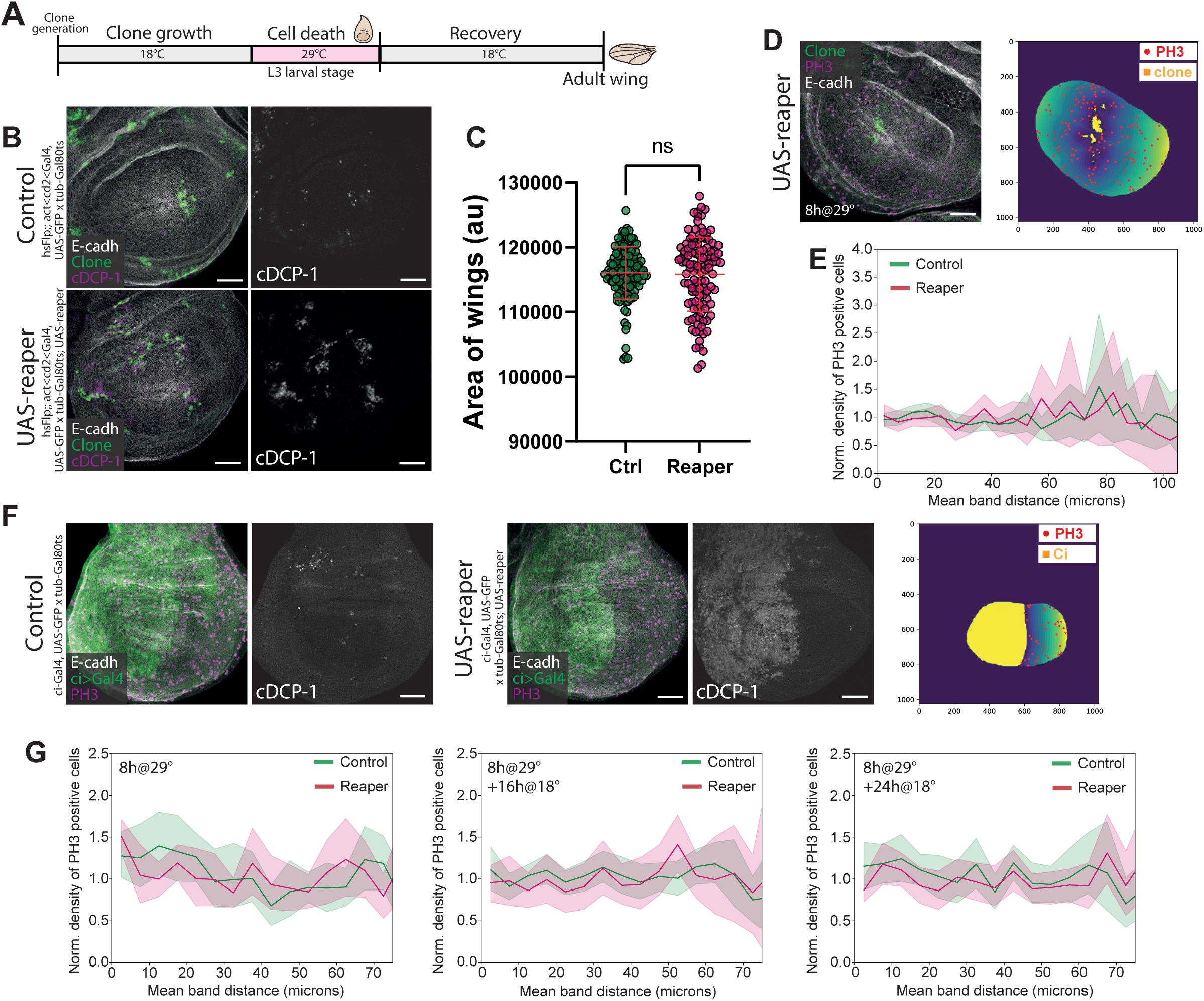
Local apoptosis does not trigger a local increase of proliferation. (**A**) Scheme of the experimental procedure. Clones are generated by a heat shock and allowed to grow at 18°C for 4 days, apoptosis through UAS-reaper (or GFP for control) is then induced by a temperature shift to 29°C (allowing the release of repression by tub-Gal80^ts^) at mid-L3 stage for 8 or 16h. Larvae are either dissected right away, or let to recover until adult stage. (**B**) Clones in control (UAS-GFP, green; act<cd2<Gal4, top) or apoptosis-induced (UAS-GFP, green ; act<cd2<Gal4 x UAS-reaper, down) wing discs after 8h at 29°C. Cleaved DCP-1 (cDCP-1, white), the *Drosophila* effector caspase, stains apoptotic cells. Scale bars, 50µm. (**C**) Size of adult wings (arbitrary units, a.u.) after apoptosis induction in clones for 16h at 96h after egg laying (AEL) (equivalent at 25°C). Ctrl (n= 128 wings, 2 experiments), Reaper (n= 115 wings, 2 experiments). Welch’s t-test, p=0.7847 (**D**) Spatial analysis of the distribution of proliferating cells after apoptosis induction in clones for 8h. Left: Proliferating cells stained by Phospho-histone H3, magenta. Scale bar, 50µm. Right: Concentric bands of 5µm (from dark blue to yellow) are drawn around clones (yellow) and the density of proliferating cells (red dots) in each band is computed by a custom-made routine in Python. (**E**) Normalised density of PH3-positive cells as a function of the distance of bands, for control (green) or apoptotic/Reaper (magenta) clones, after 8h of apoptosis induction. n=28 discs for Ctrl and n=18 disc for Reaper, from 3 independent experiments. Mean +/− 95% confidence interval. (**F**) Left: Control wing discs with GFP expressed in the anterior domain (ci-Gal4, UAS-GFP, green), showing little apoptotic cells (cDCP1, white). Middle: Apoptosis is induced for 8h in the anterior domain (ci-Gal4, UAS-GFP x UAS-reaper, green), which undergoes apoptosis (cDCP1, white). PH3 staining is shown in magenta. Scale bars, 50µm. Right: Concentric bands of 5µm in the posterior domain are drawn, starting at the boundary between the anterior (yellow) and posterior domain. The density of proliferating cells (red dots) in each band is computed by a custom-made routine in Python. (**G**) Normalised density of PH3-positive cells as a function of the distance of bands, for control (green) or apoptotic/Reaper (magenta) discs, after 8h of death induction in the anterior domain (left), followed by a recovery time of 16h (middle) or 24h (right) at 18°C. n=17 discs (ctrl 8h@29°C), n=17 discs (Reaper 8h@29°C); n=24 discs (ctrl 8h@29°C+16h@18°C), n=14 discs (Reaper 8h@29°C+16h@18°C); n=20 discs (ctrl 8h@29°C+24h@18°C), n=19 discs (Reaper 8h@29°C+24h@18°C), from 2 independent experiments. Mean +/− 95% confidence interval. See also Supplementary figures S1, S2 and S3.

### Apoptosis induction causes a transient delay of disc growth

We therefore asked which alternative mechanism could explain the compensation in the adult wing. We first checked whether apoptosis was associated with a global increase of the proliferative rate. Our first analysis revealed an apparent increase of the mitotic index soon after the release of apoptosis induction after 8 or 16h of reaper induction (**Fig. 2A**). We noticed however that all apoptotic discs appeared significantly smaller relative to control discs (−31%) right after the induction of apoptosis either in the context of clonal induction of reaper for 16h (**Fig. 2B, Fig. S4B**) or 8h (**Fig. S4A**), or upon induction of apoptosis in one compartment (**Fig. S4D**), while their size was similar prior to the activation of reaper at 29° (**Fig. S4C**) or in the absence of clone induction (**Fig. S4E,F**). This reduction cannot be explained by the removal of cells by apoptosis (which covers on average 5% of the disc and a maximum of 12%, **Fig. S1A**), and may indicate a global slow-down of growth/proliferation and/or developmental progression triggered by apoptosis induction. We could not detect a significant delay in the appearance of *achaete* expression in the dorso-ventral margin (normally appearing around 105 hours after egg laying, at mid late L3^27,28^) between the apoptotic and control discs (**Fig. 2D, E**), suggesting that the growth delay is partially uncoupled with patterning progression, as previously observed upon the perturbation of the steroid hormone Ecdysone^28^.

**Figure 2:**
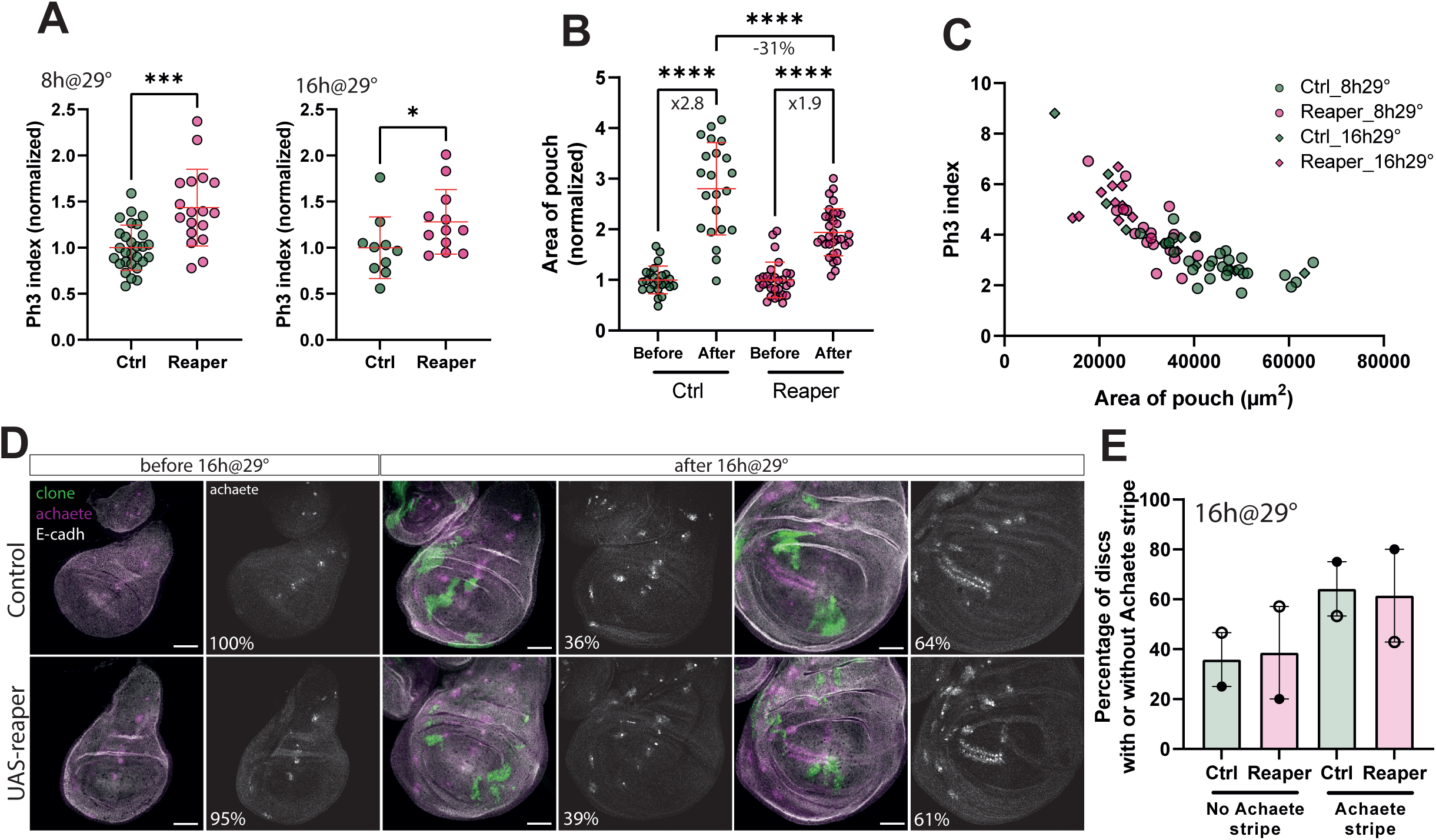
Apoptosis induction causes a transient delay of disc growth. **(A)** PH3 index in Control (green) and Reaper (magenta) conditions 8h (left) or 16h (right) after apoptosis induction in clones. PH3 index represents the number of PH3-positive cells divided by the area of the pouch. 8h@29°C: n=28 discs (Ctrl) and n=18 disc (Reaper), from 3 independent experiments. 16h@29°C: n=10 discs for (Ctrl) and n=12 discs (Reaper), in 2 independent experiments. Mann-Whitney test, p<0.05 *, p<0.01 **, p<0.001 *** **(B)** Area of the pouch at 96h AEL (before 29°C), and after 16h@29°C for Control (green) and Reaper (magenta) discs, normalised to the area at 96h AEL. n=24 (Ctrl, before), n=22 (Ctrl, after), n=29 (Reaper, before), n=32 (Reaper, after), from 2 independent experiments. One-way ANOVA, p<0.0001 ****. **(C)** PH3 index as a function of the area of the pouch for Control (green, 8h: circles, 16h: diamonds) and Reaper (magenta, 8h: circles, 16h: diamonds). n=28 (Ctrl, 8h, 3 independent experiments), n=18 (Reaper, 8h, 3 independent experiments), n=10 (Ctrl, 16h, 2 independent experiments), n=12 (Reaper, 16h, 2 independent experiments). **(D)** Wing discs from Control (top) and Reaper (bottom) larvae, at 96h AEL (before 29°C), and after 16h@29°C. Clones (green), E-cadh (white), Achaete (magenta or white). Percentages of discs with or without Achaete stripe. Scale bars, 50µm. **(E)** Percentages of discs with or without Achaete stripe, after 16h@29°C, in two different experiments (same individuals as in Figure 2B) See also Supplementary Figure S4.

The proliferation rate of the wing disc is progressively going down during the L3 stage^29^ as disc size is progressively increasing^30^. Accordingly, we found a clear negative correlation between wing pouch size and the proliferation index in the control discs (**Fig. 2C**). Strikingly, the relationship between pouch size and proliferation index is not modified in the apoptotic discs, and the mitotic index appeared unaltered when comparing disc of similar size (**Fig. 2C**, **Fig S4G**). This indicates that the apparent increase of proliferation (**Fig. 2A**) may just reflect the higher proportion of smaller/younger disc upon apoptosis induction (**Fig. 2B,C**) and is unlikely to reflect an active induction of proliferation by apoptosis.

### JNK activation in apoptotic cells and pupariation delay is required for size compensation

Induction of massive cell death in the wing disc triggers a lengthening of larval stage and delays pupariation^31,32^. Despite the mild induction of apoptosis, we also observed a slight, albeit significant, delay of pupariation time upon induction of apoptosis in clones (**Fig. 3A**, ∼4-5 hours delay) while no difference was observed between WT and UAS-reaper disc without incubation at 29° (**Fig. S5A**). We assumed that this lengthening of the larval phase may contribute to the compensation for cell loss in the wing. Apoptosis is associated with a cell autonomous induction of the stress pathway JNK^33–36^, which can delay developmental progression through the induction of the Relaxin hormone Dilp8^32,37^ and the transient inhibition of Ecdysone production^38–40^. Using the transcriptional reporter of JNK activity TRE-dsRed^41^, we observed accordingly a significant upregulation of JNK pathway in the apoptotic clones (**Fig. 3B**). We therefore checked whether JNK activation in apoptotic clones could be responsible for the non-cell autonomous slow-down of growth and the delay of the pupariation period. Transient inhibition of JNK specifically in the apoptotic clones using a dominant negative of Basket (UAS-bsk^DN^) was sufficient to abolish the wing disc growth delay observed upon apoptosis induction (**Fig. 3C**) as well as the pupariation delay induced by cell death (**Fig. 3D**). Importantly, the inhibition of JNK didn’t alter the level of apoptosis induction in the clones (**Fig S5C**). Strikingly, the adult wings could not recover their control size upon transient inhibition of JNK in apoptotic clones (**Fig. 3E**), suggesting that the compensatory mechanism is indeed driven by JNK activation in apoptotic cells and is likely to rely on the lengthening of larval stage. Developmental delay was previously shown to rely on the induction of the relaxin hormone Dilp8 downstream of JNK^32,39^. While we could not detect any clear increase of Dilp8 using LacZ or GFP reporter, RT-PCR reveals a slight increase of Dilp8 expression in apoptotic wing disc (∼90% increase, **Fig. S5B**), which may relate to the very mild level of induction of apoptosis used in this study compared to the previous characterisations of Dilp8^32^. Interestingly, we didn’t detect any global or local upregulation of the JAK-STAT^17^ or downregulation of Hippo pathways in contexts of mild induction of cell death (**Fig. S5D,E**).

**Figure 3:**
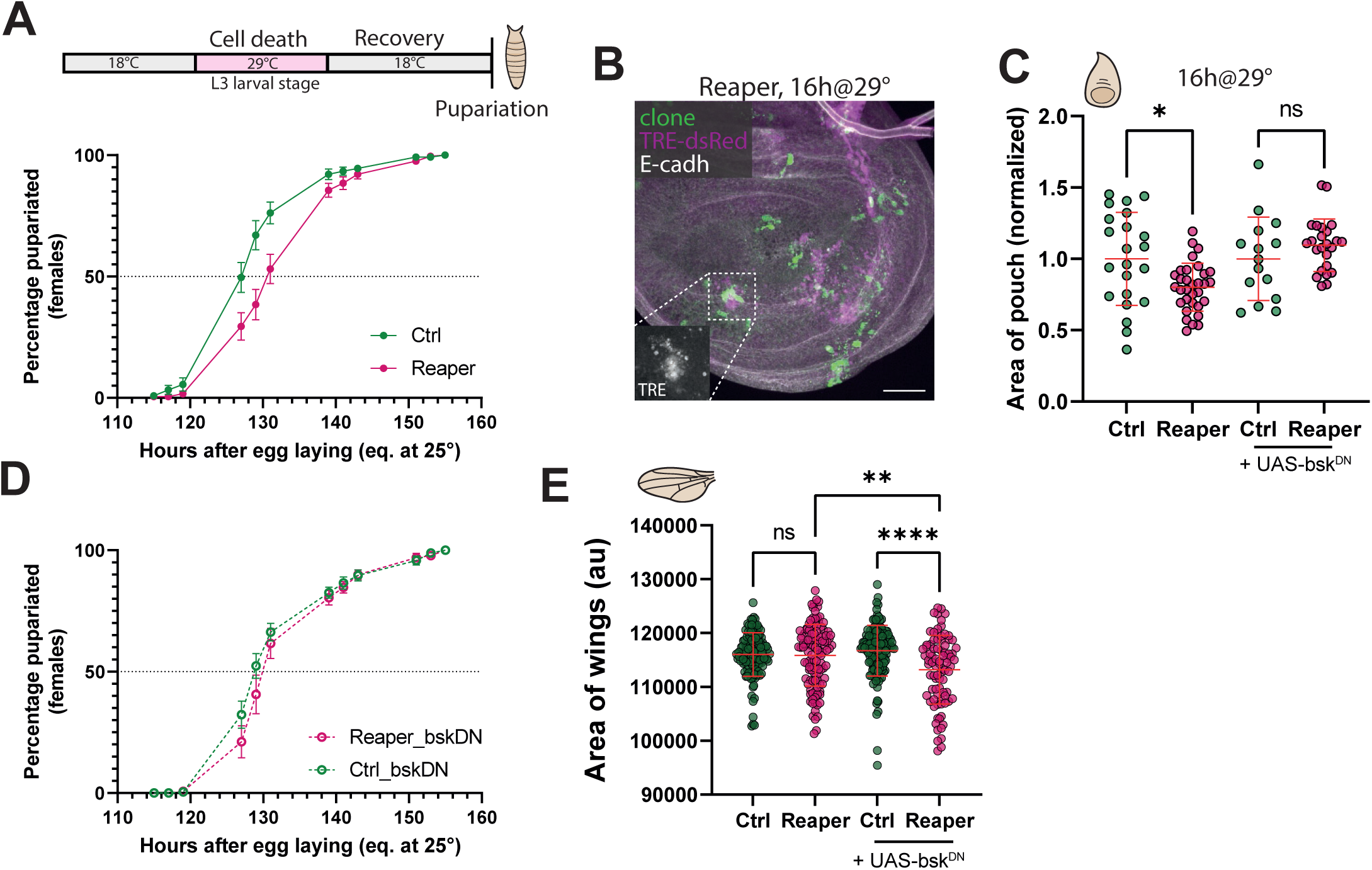
JNK activation in apoptotic cells and pupariation delay is required for size compensation. **(A)** Top: scheme of the pupariation assay. Bottom: Percentage of pupariated females (only females have both UAS-reaper and hs-flp) for Control (green) and Reaper (magenta) individuals. Clones are induced at 48h AEL and larvae are placed at 29°C for 16h at 96h AEL. n=145 females (Control) and n= 235 females (Reaper) from 2 independent experiments. Mean + SEM **(B)** TRE-RFP (magenta) expression, a JNK sensor, upon induction of apoptosis in clones. Scale bars, 50µm. **(C)** The reduction of the area of the pouch is rescued upon transient JNK inhibition in clones. n= 22 (Ctrl), n=32 (Reaper), 2 independent experiments, n=15 (Ctrl + UAS-bsk^DN^), n=24 (Reaper + UAS-bsk^DN^), one experiment. One-way ANOVA, p<0.05 *. **(D)** Percentage of pupariated females for Control (green) and Reaper (magenta) individuals upon transient JNK inhibition in clones (UAS-bsk^DN^). Clones are induced at 48h AEL and larvae are placed at 29°C for 16h at 96h AEL. n=217 females (Control) and n=123 females (Reaper) from 2 independent experiments. Mean + SEM. **(E)** Size of adult wings (arbitrary units, a.u.) after apoptosis induction in clones for 16h at 96h AEL. Ctrl (n= 128 wings, 2 experiments), Reaper (n= 115 wings, 2 experiments), Ctrl + UAS-bsk^DN^ (n= 136 wings, 2 experiments), Reaper + UAS-bsk^DN^ (n= 85 wings, 2 experiments). One-way ANOVA, p<0.01 **, p<0.0001 ****. See also Supplementary Figure S5.

### Development slow-down ensures global tissue size robustness but does not correct local alteration of growth

We reasoned that a compensatory mechanism based on global modulation of growth may be relatively inefficient for correcting local size defects (relative to a compensation driven by local induction of proliferation). This is especially true if local apoptosis triggers a global transient decrease of proliferation/growth followed by a lengthening of the growth phase that equally affects all regions of the disc. Accordingly, while adult total wing size was not reduced by induction of apoptosis in the anterior wing disc compartment (**Fig. 4A,B** the wings even appear slightly bigger), the compensation fails to correct the relative reduction of the anterior compartment size, which proportion remains smaller compared to control wings (**Fig. 4D**). These results suggest that the non-apoptotic and apoptotic regions benefit equally from the compensatory growth (**Fig. 4C**), preventing the anterior compartment from catching up with the posterior compartment. Accordingly, a similar relative decrease of the anterior compartment size is already observed at the larval stage and seems to perdure throughout development (**Fig. S6**). Moreover, transient inhibition of JNK by expressing bsk^DN^ in the anterior compartment during the apoptosis induction phase significantly reduced the absolute size of adult apoptotic wings (**Fig. 4F**, compared with reaper without Bsk^DN^), and reduced both the anterior and posterior compartment growth (**Fig. 4G,I**). As such, the A/P ratio is not altered by JNK inhibition and remains significantly smaller than the non-apoptotic control wings (**Fig. 4H,J**). These results confirmed a scenario where reduction of the anterior compartment size by a pulse of apoptosis is followed by a global growth compensation driven by JNK activity in the dying cells, which is not sufficient for the anterior compartment to catch up with the posterior one (**Fig. 4K**).

**Figure 4:**
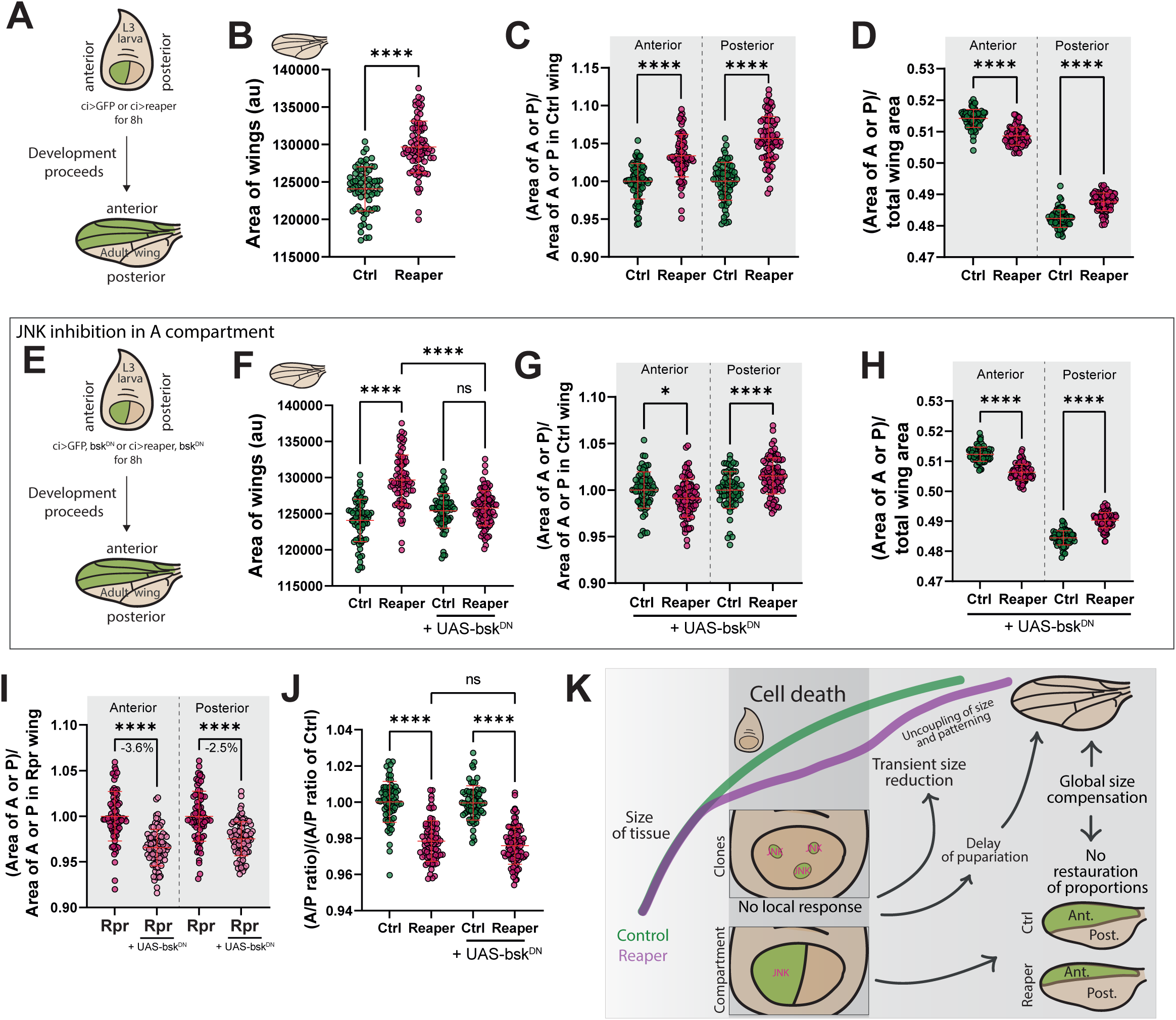
Development slow-down ensures global tissue size robustness but does not correct local alteration of growth. (**A**) Scheme of the experimental procedure. Cell death is induced in the anterior compartment for 8h at 160h AEL at 18°C (equivalent to 80h AEL at 25°C) during larval stage, and the adult wing size and proportions are assessed (**B**) Size of adult wings (arbitrary units, au) after apoptosis induction in the anterior (A) compartment. Ctrl (n= 70 wings, 1 experiment), Reaper (n=77 wings, 1 experiment). Mean + SD, Welch’s t-test, p<0.0001 ****. (**C**) Area of the anterior (A) or posterior (P) compartment normalized to the area of the corresponding compartment in Ctrl wings. Ctrl (n= 70 wings, 1 experiment), Reaper (n=77 wings, 1 experiment). Mean + SD, one-way ANOVA, p<0.0001 ****. (**D**) Area of the A or P compartment divided by the total wing area. Ctrl (n= 70 wings, 1 experiment), Reaper (n=77 wings, 1 experiment). Mean + SD, one-way ANOVA, p<0.0001****. (**E**) Cell death and concomitant JNK inhibition (UAS-bsk^DN^) are induced in the anterior compartment for 8h at 160h AEL at 18°C (equivalent to 80h AEL at 25°C) during larval stage, and the adult wing size and proportions are assessed. (**F**) Size of adult wings (arbitrary units, a.u.) after apoptosis induction in the anterior (A) compartment. Ctrl (n= 70 wings, 1 experiment), Reaper (n=77 wings, 1 experiment), Ctrl+UAS-bsk^DN^ (n= 69 wings, 1 experiment), Reaper+UAS-bsk^DN^ (n=91 wings, 1 experiment). Mean + SD, one-way ANOVA, p<0.0001 ****. (**G**) Area of the anterior (A) or posterior (P) compartment normalized to the area of the corresponding compartment in Ctrl wings. Ctrl+UAS-bsk^DN^ (n= 69 wings, 1 experiment), Reaper+UAS-bsk^DN^ (n=91 wings, 1 experiment). Mean + SD, one-way ANOVA, p<0.05 *, p<0.0001 ****. (**H**) Area of the A or P compartment divided by the total wing area. Ctrl+UAS-bsk^DN^ (n= 69 wings, 1 experiment), Reaper+UAS-bsk^DN^ (n=91 wings, 1 experiment). Mean + SD, one-way ANOVA, p<0.0001 ****. (**I**) Area of the A or P compartment divided by the area of the corresponding A or P compartment in the Reaper wing. Reaper (n= 77 wings, 1 experiment), Reaper+UAS-bsk^DN^ (n=91 wings, 1 experiment). Mean + SD, one-way ANOVA, p<0.0001 (**J**) A/P ratio divided by the A/P of Ctrl. Ctrl (n= 70 wings, 1 experiment), Reaper (n=77 wings, 1 experiment), Ctrl+UAS-bsk^DN^ (n= 69 wings, 1 experiment), Reaper+UAS-bsk^DN^ (n=91 wings, 1 experiment). Mean + SD, one-way ANOVA, p<0.0001 ****. (**K**) Scheme of the proposed mechanism. See also Supplementary Figure S6.

## Discussion

Here we provided one of the first quantitative dissections of the reaction of the wing disc to a mild induction of apoptosis. We first confirmed that the adult wing could indeed compensate for the loss of 4 to 12% of cells, revealing once again the striking robustness of size regulation. Surprisingly, we could not find any sign of local induction of proliferation (**Fig. 1**, **Fig. S1**, **S2 and S3**) and rather observed a transient global decrease of growth followed by a lengthening of the larval period (**Fig. 2**) driven by transient JNK activation in the apoptotic cells (**Fig. 3**). Interestingly, a similar developmental delay-based mechanism was recently reported upon global disc irradiation^42^. Using systematic quantitative mapping of cell death and clone disappearance, we recently outlined the existence of apoptosis hotspots in the wing disc which are associated with a local reduction of net clone size and absolute growth^2^. This was already suggesting that physiological cell death was not necessarily locally compensated by more proliferation/growth, in good agreement with the absence of significant response observed in this study upon mild ectopic apoptosis induction. These results contrast with the large body of literature reporting local upregulation of cell proliferation upon damage and suggest that the compensatory response may be highly context dependent and may vary accordingly to the level and duration of apoptotic induction. Compensatory proliferation driven by undead cells relies on the accumulation of the active Caspase 9 Dronc which triggers JNK activation and secretion of various mitogens such as Dpp, Wg or EGF^34–36^. Such response should rely on the relatively long perdurance of cells with active Dronc which is unlikely to happen in the context of rapid apoptotic cell clearance. Accordingly, these local signals were confirmed in the context of physiological gut enterocyte apoptosis^43,44^, where single cell apoptosis and cell extrusion takes place over several hours^45^. Alternatively, the high rate of apoptosis induced in a compartment for long periods of time or upon irradiation may alter core tissue properties that become permissive for local signal propagation and compensatory growth. For instance, high apoptosis rate may affect epithelial sealing properties^25,46^ hence modulating signal propagation and the accessibility to some ligands^47^. Alternatively, high apoptosis rates may modulate significantly the global mechanical properties of the tissue which can fine tune the responsiveness of neighbouring cells to apoptosis, as illustrated in MDCK cells^23^. Further quantitative work would help to define the minimal conditions allowing the observation of such local growth compensation.

Surprisingly, we also found that the phase of apoptosis induction correlates with a transient reduction of disc size, most likely through a tissue wide slow-down of cell growth and cell division. As such, apoptotic discs have to compensate both for the loss of cells driven by apoptosis and the global growth delay associated with apoptosis induction. The secretion of Dilp8 from damaged tissue was proposed to contribute to tissue regeneration through the lengthening of larval phase driven by Ecdysone downregulation^32,37^. Paradoxically, basal Ecdysone levels also have an important pro-proliferative/growth role^30,48,49^ and the downregulation of Ecdysone should also slow-down tissue size recovery, as observed in our tissues during the apoptosis-induction phase. While the lengthening of larval stage alone may be sufficient to compensate tissue damage and growth slow-down, other mechanisms may also contribute to the final size recovery. For instance, an increased sensitivity to Ecdysone levels after size reduction and/or a rapid induction of growth driven by Ecdysone concentration increase over time could help to accelerate disc growth. Accordingly, smaller discs were shown to be more responsive to Ecdysone relative to larger discs^30^, which could explain the higher proliferative index observed in the smaller apoptotic discs after the apoptotic phase (**Fig. 2A,C**). This response may be amplified by a non-linear response to Ecdysone concentrations^28,50^. Interestingly, our results also suggest that the delay in growth triggered by JNK and apoptosis is uncoupled with the progression of patterning gene expression (**Fig. 2D,E**). This is in good agreement with previous results showing linear response of growth to Ecdysone, and a logistic/threshold control for patterning genes^28^, suggesting that Ecdysone would pass the *achaete* induction threshold at the same time in Reaper and control discs, while growth would be altered by the transient Ecdysone reduction.

The most striking conclusion of our work is that the recovery from local apoptosis mostly relies on a global tissue level response rather than a local induction of proliferation. This spatial difference should have important consequences for size regulation and robustness. Accordingly, we found that the relative decrease of a compartment size driven by transient induction of apoptosis could not be totally rescued in the adult wing, while global wing size could be (**Fig. 4**). These results contrast with previous works showing that long term alteration of growth in one compartment slows-down growth in the adjacent compartment to maintain proportions^51,52^. This could reflect a difference between a transient induction of apoptosis, where global growth slow-down is followed by global compensation, and a long term/chronic stress induction where the two phases are intermingled. Further theoretical work may be required to understand whether such integral control rather than local compensation carries some advantages for the robustness of tissue morphogenesis, or whether this a simple constrain of the developmental hardwiring of growth regulation and evolutionary “bricolage/tinkering”. Cell competition is an ubiquitous mechanism that leads to the context-dependant elimination of one cell population when interacting with another more competitive cell population^53–55^. This mechanism is thought to promote the expansion of the so-called winner cells through the induction of apoptosis in the loser population and the subsequent local compensatory proliferation of winner cells located near loser cells, hence leaving tissue size unaffected^56^. However, a global response to loser cell death, as suggested by our study, would dramatically change the consequences of compensatory growth on the outcome of cell competition and make the specific localisation of cell death in loser clones quite irrelevant to the outcome of cell competition. Further quantitative characterisation of the spatio-temporal pattern of cell growth and cell proliferation in cell competition contexts would be essential to fully understand the contribution of compensatory growth for the final outcome of competition.

## Acknowledgements

We would like thank members of the CDEH group for constructive comments on the manuscript. We are also grateful to Pierre Leopold, Laura Boulan, Dalmiro Blanco Obregon, Marco Milan, the Bloomington Drosophila Stock Center, the Drosophila Genetic Resource Center and the Vienna Drosophila Resource Center, Flybase for sharing essential information, and the Developmental Studies Hybridoma Bank for stocks and reagents. We also thank the image analysis hub (IAH) of Institut Pasteur for generating a Python code for spatial analysis (Marvin Albert). Work in RL lab is supported by the Institut Pasteur, the European Union (ERC, PrApEDoC, 101085444), Views and opinions expressed are however those of the author(s) only and do not necessarily reflect those of the European Union or the European Research Council. Neither the European Union nor the granting authority can be held responsible for them, the ANR PRC CoECECa, the ANR PRC MAPEFLU, and the ANR-10-LABX-0073 and the CNRS (UMR 3738). Ralitza Staneva was supported by an ARC postdoctoral grant Aide individuelle (PDF20191209565), and a Pasteur Roux-Cantarini fellowship.

## Authors contributions

R.S. and R.L. designed and discussed the project and wrote the manuscript. G.S.M and A.D. performed the K and G function analysis, A.D. supervised G.S.M.. F.L. performed the QPCR. A.V. helped for the analysis of spatial distribution and randomisation. R.S. performed all the other experiments and analysis. All the authors commented and edited the manuscript.

## Star Methods

### Ressource availability

#### Lead contact

Further information and requests for resources and reagents should be directed to and will be fulfilled by the lead contact, Romain Levayer.

#### Material availability

All the reagents generated in this study will be shared upon request to the lead contact without any restrictions.

#### Data and Code availability

All code generated in this study and the raw data corresponding to each figure panel will be accessible on a repository upon final publication of the article. Further information about the dataset can be requested to the lead contact.

### Experimental model and subject details

#### Drosophila melanogaster husbandry

All the experiments were performed with *Drosophila melanogaster* fly lines with regular husbandry technics. The fly food used contains agar agar (7.6 g/l), saccharose (53 g/l) dry yeast (48 g/l), maize flour (38.4 g/l), propionic acid (3.8 ml/l), Nipagin 10% (23.9 ml/l) all mixed in one liter of distilled water. Flies were raised at 25°C in plastic vials with a 12h/12h dark light cycle at 60% of moisture unless specified in the legends and in the table below. We did not determine the health/immune status of embryos, larvae and adults, they were not involved in previous procedures, and they were all drug and test naïve.

#### Drosophila melanogaster strains

The strains used in this study and their origin are listed in the table below.

| Fly line | Chromosome location | Origin (citation) | RRID |
| --- | --- | --- | --- |
| *hs-flp; if/Cyo; act<cd2<Gal4, UAS-GFP/TM6b* | III | Eduardo Moreno | N/A |
| *UAS-reaper; tub-Gal80<sup>ts</sup>/Cyo* | X; II | Eduardo Moreno | N/A |
| *hs-flp22; tub-Gal80<sup>ts</sup>/Cyo* | X, II | Bloomington | BDSC_7108 |
| *w; ci-Gal4, UAS-GFP* | II | Marco Milan | N/A |
| *w1118; ; STAT-GFP* | III | Bloomington | BDSC_26198 |
| *expanded-LacZ* | II | Bloomington | BDSC_44248 |
| *TRE-dsRed-40* | II | Dirk Bohman <sup>41</sup> | N/A |
| *hs-flp22; act<y<Gal4, UAS-mcd8-RFP/Cyo; MKRS/TM6b* | X, II, III | Romain Levayer | N/A |
| *UAS-bsk<sup>DN</sup>* | III | Bloomington | BDSC_9311 |

The exact genotype used for each experiment is listed in the next table. ACI: time After Clone Induction, AEL: time After Egg Laying, n: number of wing discs/adults. Only females are used.

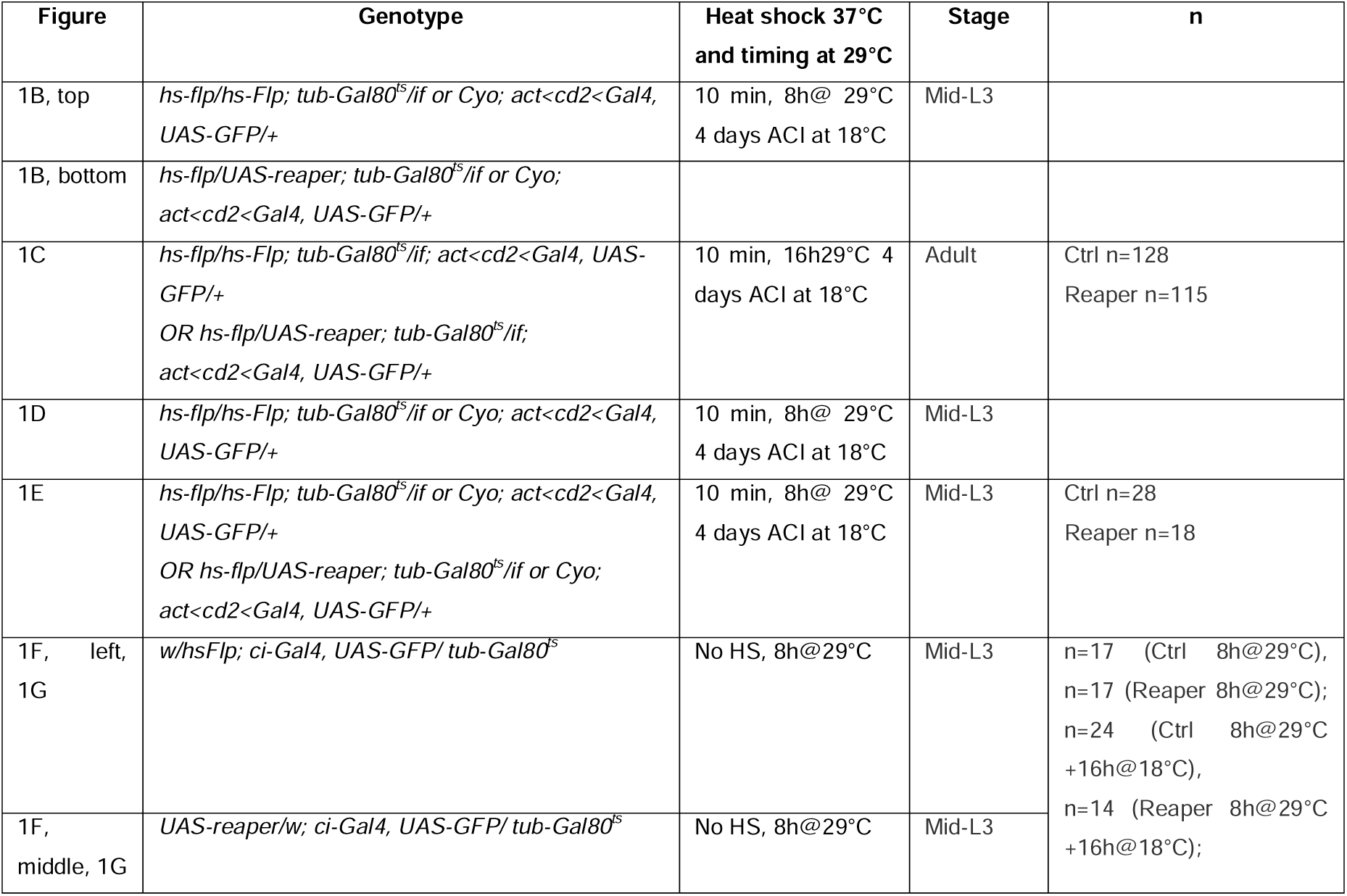
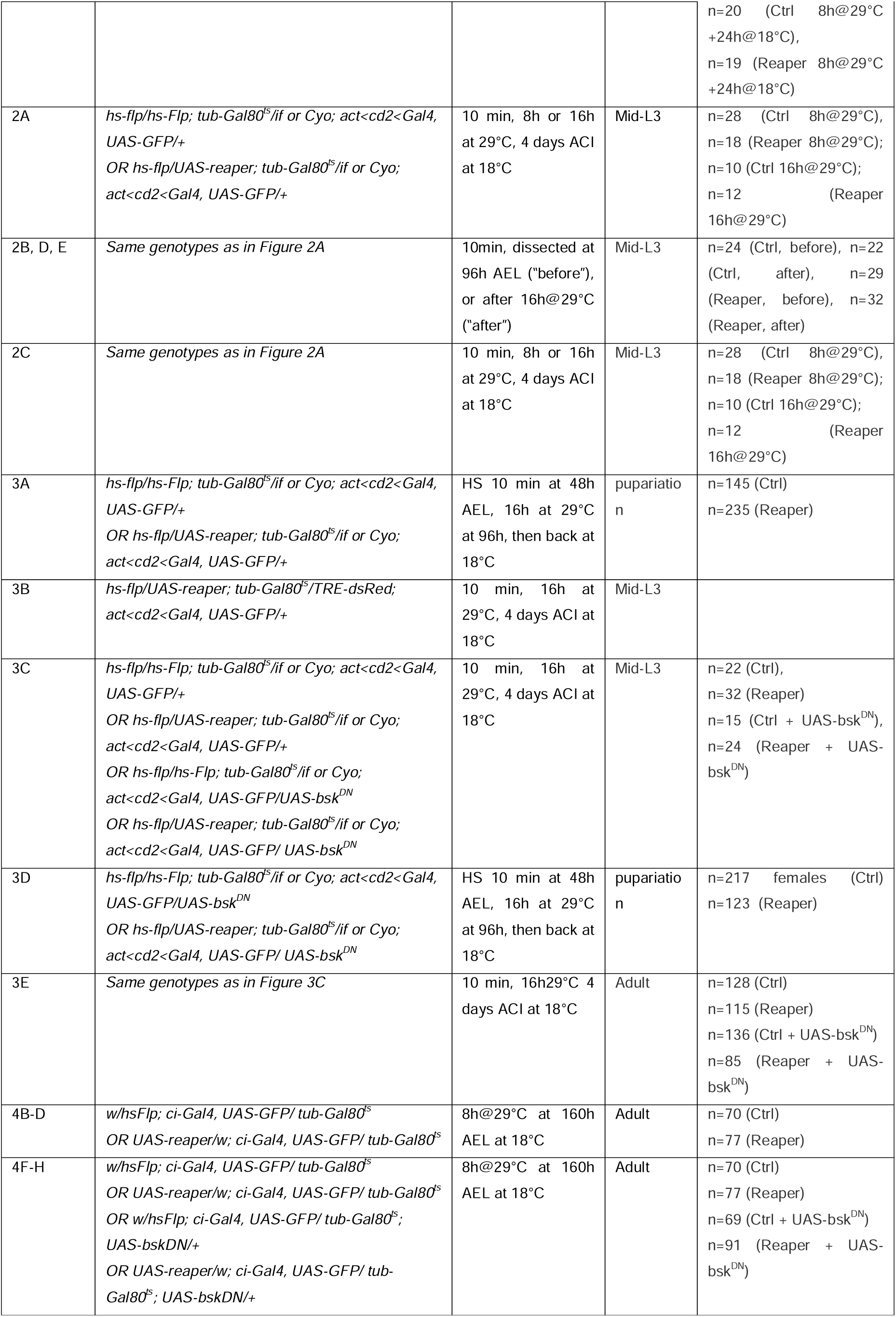
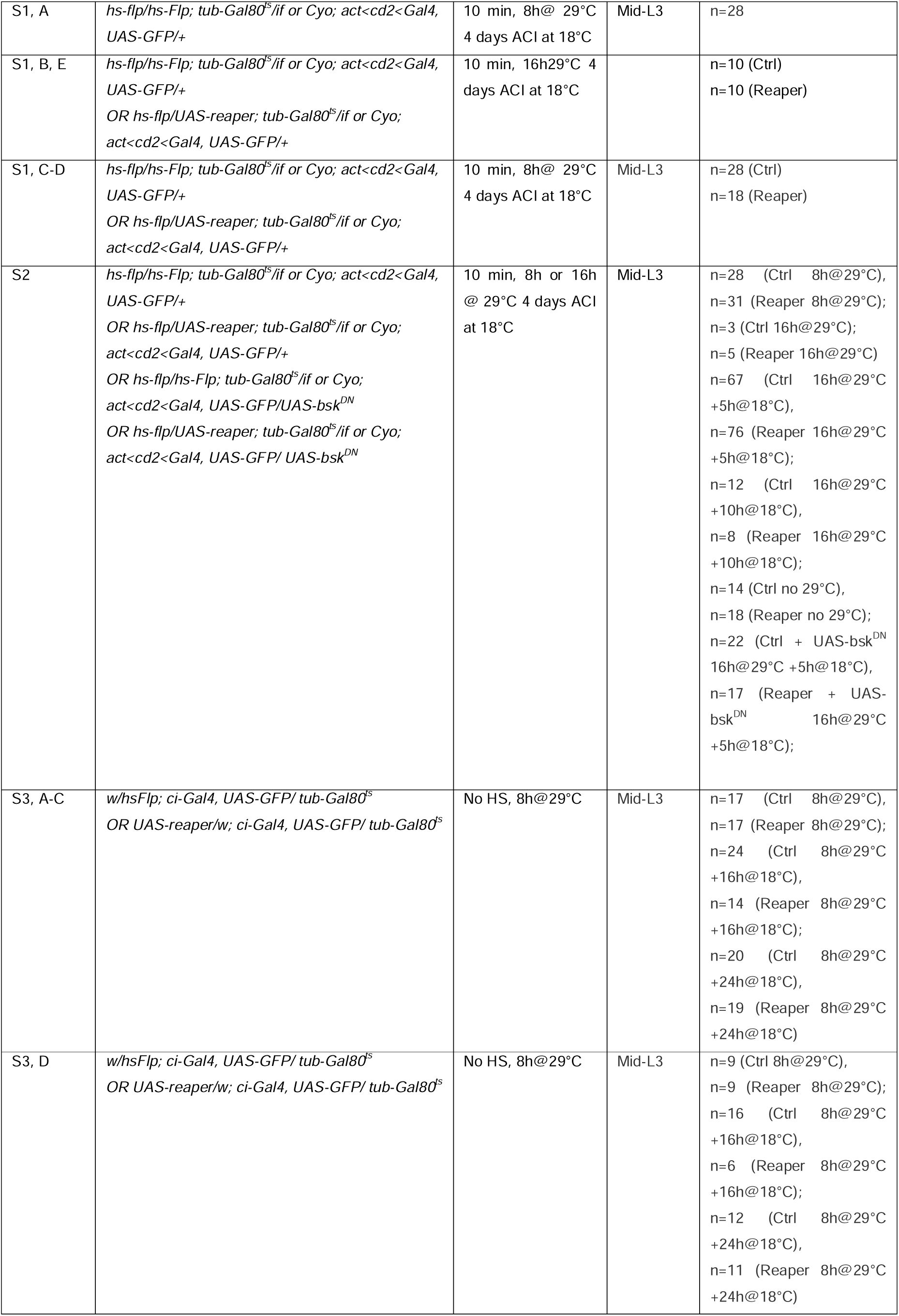
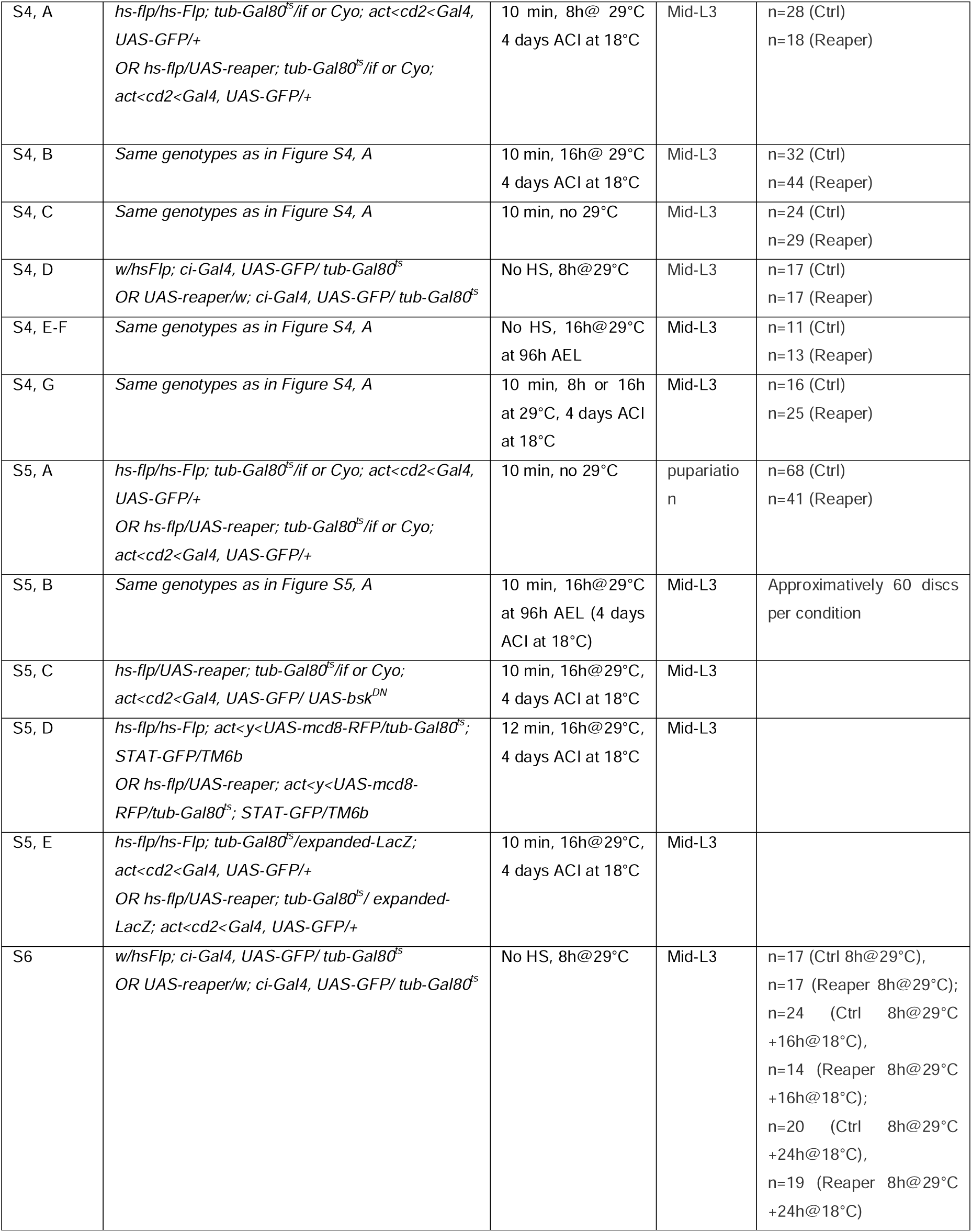

| Figure | Genotype | Heat shock 37°C and timing at 29°C | Stage | n |
| --- | --- | --- | --- | --- |
| 1B, top | *hs-flp/hs-Flp; tub-Gal80<sup>ts</sup>/if or Cyo; act<cd2<Gal4, UAS-GFP/+* | 10 min, 8h@ 29°C<br>4 days ACI at 18°C | Mid-L3 |  |
| 1B, bottom | *hs-flp/UAS-reaper; tub-Gal80<sup>ts</sup>/if or Cyo; act<cd2<Gal4, UAS-GFP/+* |  |  |  |
| 1C | *hs-flp/hs-Flp; tub-Gal80<sup>ts</sup>/if; act<cd2<Gal4, UAS-GFP/+*<br>OR *hs-flp/UAS-reaper; tub-Gal80<sup>ts</sup>/if; act<cd2<Gal4, UAS-GFP/+* | 10 min, 16h@29°C<br>4 days ACI at 18°C | Adult | Ctrl n=128<br>Reaper n=115 |
| 1D | *hs-flp/hs-Flp; tub-Gal80<sup>ts</sup>/if or Cyo; act<cd2<Gal4, UAS-GFP/+* | 10 min, 8h@ 29°C<br>4 days ACI at 18°C | Mid-L3 |  |
| 1E | *hs-flp/hs-Flp; tub-Gal80<sup>ts</sup>/if or Cyo; act<cd2<Gal4, UAS-GFP/+*<br>OR *hs-flp/UAS-reaper; tub-Gal80<sup>ts</sup>/if or Cyo; act<cd2<Gal4, UAS-GFP/+* | 10 min, 8h@ 29°C<br>4 days ACI at 18°C | Mid-L3 | Ctrl n=28<br>Reaper n=18 |
| 1F, left, 1G | *w/hsFlp; ci-Gal4, UAS-GFP/ tub-Gal80<sup>ts</sup>* | No HS, 8h@29°C | Mid-L3 | n=17 (Ctrl 8h@29°C),<br>n=17 (Reaper 8h@29°C);<br>n=24 (Ctrl 8h@29°C +16h@18°C),<br>n=14 (Reaper 8h@29°C +16h@18°C); |
| 1F, middle, 1G | *UAS-reaper/w; ci-Gal4, UAS-GFP/ tub-Gal80<sup>ts</sup>* | No HS, 8h@29°C | Mid-L3 |  |
|  |  |  |  | n=20 (Ctrl 8h@29°C +24h@18°C),<br>n=19 (Reaper 8h@29°C +24h@18°C) |
| 2A | *hs-flp/hs-Flp; tub-Gal80<sup>ts</sup>/if or Cyo; act<cd2<Gal4, UAS-GFP/+*<br>OR *hs-flp/UAS-reaper; tub-Gal80<sup>ts</sup>/if or Cyo; act<cd2<Gal4, UAS-GFP/+* | 10 min, 8h or 16h at 29°C, 4 days ACI at 18°C | Mid-L3 | n=28 (Ctrl 8h@29°C),<br>n=18 (Reaper 8h@29°C);<br>n=10 (Ctrl 16h@29°C);<br>n=12 (Reaper 16h@29°C) |
| 2B, D, E | Same genotypes as in Figure 2A | 10min, dissected at 96h AEL ("before"), or after 16h@29°C ("after") | Mid-L3 | n=24 (Ctrl, before), n=22 (Ctrl, after), n=29 (Reaper, before), n=32 (Reaper, after) |
| 2C | Same genotypes as in Figure 2A | 10 min, 8h or 16h at 29°C, 4 days ACI at 18°C | Mid-L3 | n=28 (Ctrl 8h@29°C),<br>n=18 (Reaper 8h@29°C);<br>n=10 (Ctrl 16h@29°C);<br>n=12 (Reaper 16h@29°C) |
| 3A | *hs-flp/hs-Flp; tub-Gal80<sup>ts</sup>/if or Cyo; act<cd2<Gal4, UAS-GFP/+*<br>OR *hs-flp/UAS-reaper; tub-Gal80<sup>ts</sup>/if or Cyo; act<cd2<Gal4, UAS-GFP/+* | HS 10 min at 48h AEL, 16h at 29°C at 96h, then back at 18°C | pupariation | n=145 (Ctrl)<br>n=235 (Reaper) |
| 3B | *hs-flp/UAS-reaper; tub-Gal80<sup>ts</sup>/TRE-dsRed; act<cd2<Gal4, UAS-GFP/+* | 10 min, 16h at 29°C, 4 days ACI at 18°C | Mid-L3 |  |
| 3C | *hs-flp/hs-Flp; tub-Gal80<sup>ts</sup>/if or Cyo; act<cd2<Gal4, UAS-GFP/+*<br>OR *hs-flp/UAS-reaper; tub-Gal80<sup>ts</sup>/if or Cyo; act<cd2<Gal4, UAS-GFP/+*<br>OR *hs-flp/hs-Flp; tub-Gal80<sup>ts</sup>/if or Cyo; act<cd2<Gal4, UAS-GFP/UAS-bsk<sup>DN</sup>*<br>OR *hs-flp/UAS-reaper; tub-Gal80<sup>ts</sup>/if or Cyo; act<cd2<Gal4, UAS-GFP/ UAS-bsk<sup>DN</sup>* | 10 min, 16h at 29°C, 4 days ACI at 18°C | Mid-L3 | n=22 (Ctrl),<br>n=32 (Reaper)<br>n=15 (Ctrl + UAS-bsk<sup>DN</sup>),<br>n=24 (Reaper + UAS-bsk<sup>DN</sup>) |
| 3D | *hs-flp/hs-Flp; tub-Gal80<sup>ts</sup>/if or Cyo; act<cd2<Gal4, UAS-GFP/UAS-bsk<sup>DN</sup>*<br>OR *hs-flp/UAS-reaper; tub-Gal80<sup>ts</sup>/if or Cyo; act<cd2<Gal4, UAS-GFP/ UAS-bsk<sup>DN</sup>* | HS 10 min at 48h AEL, 16h at 29°C at 96h, then back at 18°C | pupariation | n=217 females (Ctrl)<br>n=123 (Reaper) |
| 3E | Same genotypes as in Figure 3C | 10 min, 16h@29°C 4 days ACI at 18°C | Adult | n=128 (Ctrl)<br>n=115 (Reaper)<br>n=136 (Ctrl + UAS-bsk<sup>DN</sup>)<br>n=85 (Reaper + UAS-bsk<sup>DN</sup>) |
| 4B-D | *w/hsFlp; ci-Gal4, UAS-GFP/ tub-Gal80<sup>ts</sup>*<br>OR *UAS-reaper/w; ci-Gal4, UAS-GFP/ tub-Gal80<sup>ts</sup>* | 8h@29°C at 160h AEL at 18°C | Adult | n=70 (Ctrl)<br>n=77 (Reaper) |
| 4F-H | *w/hsFlp; ci-Gal4, UAS-GFP/ tub-Gal80<sup>ts</sup>*<br>OR *UAS-reaper/w; ci-Gal4, UAS-GFP/ tub-Gal80<sup>ts</sup>*<br>OR *w/hsFlp; ci-Gal4, UAS-GFP/ tub-Gal80<sup>ts</sup>; UAS-bskDN/+*<br>OR *UAS-reaper/w; ci-Gal4, UAS-GFP/ tub-Gal80<sup>ts</sup>; UAS-bskDN/+* | 8h@29°C at 160h AEL at 18°C | Adult | n=70 (Ctrl)<br>n=77 (Reaper)<br>n=69 (Ctrl + UAS-bsk<sup>DN</sup>)<br>n=91 (Reaper + UAS-bsk<sup>DN</sup>) |
| S1, A | *hs-flp/hs-Flp; tub-Gal80<sup>ts</sup>/if or Cyo; act<cd2<Gal4, UAS-GFP/+* | 10 min, 8h@ 29°C<br>4 days ACI at 18°C | Mid-L3 | n=28 |
| S1, B, E | *hs-flp/hs-Flp; tub-Gal80<sup>ts</sup>/if or Cyo; act<cd2<Gal4, UAS-GFP/+*<br>*OR hs-flp/UAS-reaper; tub-Gal80<sup>ts</sup>/if or Cyo; act<cd2<Gal4, UAS-GFP/+* | 10 min, 16h@29°C<br>4 days ACI at 18°C |  | n=10 (Ctrl)<br>n=10 (Reaper) |
| S1, C-D | *hs-flp/hs-Flp; tub-Gal80<sup>ts</sup>/if or Cyo; act<cd2<Gal4, UAS-GFP/+*<br>*OR hs-flp/UAS-reaper; tub-Gal80<sup>ts</sup>/if or Cyo; act<cd2<Gal4, UAS-GFP/+* | 10 min, 8h@ 29°C<br>4 days ACI at 18°C | Mid-L3 | n=28 (Ctrl)<br>n=18 (Reaper) |
| S2 | *hs-flp/hs-Flp; tub-Gal80<sup>ts</sup>/if or Cyo; act<cd2<Gal4, UAS-GFP/+*<br>*OR hs-flp/UAS-reaper; tub-Gal80<sup>ts</sup>/if or Cyo; act<cd2<Gal4, UAS-GFP/+*<br>*OR hs-flp/hs-Flp; tub-Gal80<sup>ts</sup>/if or Cyo; act<cd2<Gal4, UAS-GFP/UAS-bsk<sup>DN</sup>*<br>*OR hs-flp/UAS-reaper; tub-Gal80<sup>ts</sup>/if or Cyo; act<cd2<Gal4, UAS-GFP/ UAS-bsk<sup>DN</sup>* | 10 min, 8h or 16h<br>@ 29°C 4 days ACI<br>at 18°C | Mid-L3 | n=28 (Ctrl 8h@29°C),<br>n=31 (Reaper 8h@29°C);<br>n=3 (Ctrl 16h@29°C);<br>n=5 (Reaper 16h@29°C)<br>n=67 (Ctrl 16h@29°C<br>+5h@18°C),<br>n=76 (Reaper 16h@29°C<br>+5h@18°C);<br>n=12 (Ctrl 16h@29°C<br>+10h@18°C),<br>n=8 (Reaper 16h@29°C<br>+10h@18°C);<br>n=14 (Ctrl no 29°C),<br>n=18 (Reaper no 29°C);<br>n=22 (Ctrl + UAS-bsk<sup>DN</sup><br>16h@29°C +5h@18°C),<br>n=17 (Reaper + UAS-bsk<sup>DN</sup> 16h@29°C<br>+5h@18°C); |
| S3, A-C | *w/hsFlp; ci-Gal4, UAS-GFP/ tub-Gal80<sup>ts</sup>*<br>*OR UAS-reaper/w; ci-Gal4, UAS-GFP/ tub-Gal80<sup>ts</sup>* | No HS, 8h@29°C | Mid-L3 | n=17 (Ctrl 8h@29°C),<br>n=17 (Reaper 8h@29°C);<br>n=24 (Ctrl 8h@29°C<br>+16h@18°C),<br>n=14 (Reaper 8h@29°C<br>+16h@18°C);<br>n=20 (Ctrl 8h@29°C<br>+24h@18°C),<br>n=19 (Reaper 8h@29°C<br>+24h@18°C) |
| S3, D | *w/hsFlp; ci-Gal4, UAS-GFP/ tub-Gal80<sup>ts</sup>*<br>*OR UAS-reaper/w; ci-Gal4, UAS-GFP/ tub-Gal80<sup>ts</sup>* | No HS, 8h@29°C | Mid-L3 | n=9 (Ctrl 8h@29°C),<br>n=9 (Reaper 8h@29°C);<br>n=16 (Ctrl 8h@29°C<br>+16h@18°C),<br>n=6 (Reaper 8h@29°C<br>+16h@18°C);<br>n=12 (Ctrl 8h@29°C<br>+24h@18°C),<br>n=11 (Reaper 8h@29°C<br>+24h@18°C) |
| S4, A | *hs-flp/hs-Flp; tub-Gal80<sup>ts</sup>/if or Cyo; act<cd2<Gal4, UAS-GFP/+*<br>OR *hs-flp/UAS-reaper; tub-Gal80<sup>ts</sup>/if or Cyo; act<cd2<Gal4, UAS-GFP/+* | 10 min, 8h@ 29°C<br>4 days ACI at 18°C | Mid-L3 | n=28 (Ctrl)<br>n=18 (Reaper) |
| S4, B | Same genotypes as in Figure S4, A | 10 min, 16h@ 29°C<br>4 days ACI at 18°C | Mid-L3 | n=32 (Ctrl)<br>n=44 (Reaper) |
| S4, C | Same genotypes as in Figure S4, A | 10 min, no 29°C | Mid-L3 | n=24 (Ctrl)<br>n=29 (Reaper) |
| S4, D | *w/hsFlp; ci-Gal4, UAS-GFP/ tub-Gal80<sup>ts</sup>*<br>OR *UAS-reaper/w; ci-Gal4, UAS-GFP/ tub-Gal80<sup>ts</sup>* | No HS, 8h@29°C | Mid-L3 | n=17 (Ctrl)<br>n=17 (Reaper) |
| S4, E-F | Same genotypes as in Figure S4, A | No HS, 16h@29°C<br>at 96h AEL | Mid-L3 | n=11 (Ctrl)<br>n=13 (Reaper) |
| S4, G | Same genotypes as in Figure S4, A | 10 min, 8h or 16h<br>at 29°C, 4 days ACI<br>at 18°C | Mid-L3 | n=16 (Ctrl)<br>n=25 (Reaper) |
| S5, A | *hs-flp/hs-Flp; tub-Gal80<sup>ts</sup>/if or Cyo; act<cd2<Gal4, UAS-GFP/+*<br>OR *hs-flp/UAS-reaper; tub-Gal80<sup>ts</sup>/if or Cyo; act<cd2<Gal4, UAS-GFP/+* | 10 min, no 29°C | pupariation | n=68 (Ctrl)<br>n=41 (Reaper) |
| S5, B | Same genotypes as in Figure S5, A | 10 min, 16h@29°C<br>at 96h AEL (4 days<br>ACI at 18°C) | Mid-L3 | Approximately 60 discs<br>per condition |
| S5, C | *hs-flp/UAS-reaper; tub-Gal80<sup>ts</sup>/if or Cyo; act<cd2<Gal4, UAS-GFP/ UAS-bsk<sup>DN</sup>* | 10 min, 16h@29°C,<br>4 days ACI at 18°C | Mid-L3 |  |
| S5, D | *hs-flp/hs-Flp; act<y<UAS-mcd8-RFP/tub-Gal80<sup>ts</sup>; STAT-GFP/TM6b*<br>OR *hs-flp/UAS-reaper; act<y<UAS-mcd8-RFP/tub-Gal80<sup>ts</sup>; STAT-GFP/TM6b* | 12 min, 16h@29°C,<br>4 days ACI at 18°C | Mid-L3 |  |
| S5, E | *hs-flp/hs-Flp; tub-Gal80<sup>ts</sup>/expanded-LacZ; act<cd2<Gal4, UAS-GFP/+*<br>OR *hs-flp/UAS-reaper; tub-Gal80<sup>ts</sup>/ expanded-LacZ; act<cd2<Gal4, UAS-GFP/+* | 10 min, 16h@29°C,<br>4 days ACI at 18°C | Mid-L3 |  |
| S6 | *w/hsFlp; ci-Gal4, UAS-GFP/ tub-Gal80<sup>ts</sup>*<br>OR *UAS-reaper/w; ci-Gal4, UAS-GFP/ tub-Gal80<sup>ts</sup>* | No HS, 8h@29°C | Mid-L3 | n=17 (Ctrl 8h@29°C),<br>n=17 (Reaper 8h@29°C);<br>n=24 (Ctrl 8h@29°C<br>+16h@18°C),<br>n=14 (Reaper 8h@29°C<br>+16h@18°C);<br>n=20 (Ctrl 8h@29°C<br>+24h@18°C),<br>n=19 (Reaper 8h@29°C<br>+24h@18°C) |

### qRT-PCR for Dilp8

Total RNA was prepared from approximately 20 wing imaginal discs per condition using the QIAGEN RNeasy Plus Mini Kit. RNA was retrotranscribed using random primers and Superscript III reverse transcriptase (Invitrogen). cDNA was analyzed by qPCR using FastStart Universal SYBR Green (Roche). Actin-42A was used as a reference gene. All assays were performed in quadruplicates, and mean values were calculated according to the ΔΔCT quantification method.

Actin 42A:

Sense primer: 5’-GAGCGCGGTTACAGCTTCA-3’

Antisense primer: 5’-TCCTTGATGTCGCGCACA-3’

Dilp8:

Sense primer: 5’-CCACCATCTGAATCGACTTGG-3’

Antisense primer: 5’-CCATTGGATAGCTGCTTCGG-3’

### Wing disc dissection, immunostaining and imaging

Only female larvae were used. Wing discs were dissected in PBS on ice and fixed for 20min in 4% formaldehyde (Sigma F8775). Discs were rinsed 3 times in PBS-T (PBS + 0.3% Triton) and primary antibodies were incubated overnight at 4°C under agitation. Discs were rinsed 3 times in PBS-T and washed 3 times for a total of 2h. Secondary antibodies were incubated for 2h at room temperature, followed by 3 rinses with PBS-T and 3 washes of 30min. For achaete staining, discs were incubated in a blocking solution (PBS +1% Triton +2% normal Donkey Serum, Jackson ImmunoResearch AB_2337258) for 30min before primary antibodies incubation and PBS+1% Triton was used instead of PBS+0.3% Triton. Discs were mounted in Vectashield (Eurobio Scientific H-1000) and imaged using a laser-scanning confocal LSM880 equipped with an oil 40X objective (N.A. 1.3) with z-stacks of 0.5µm.

The following primary antibodies were used: rat anti-E-cadherin (1:100, DCAD2 concentrated DSHB), mouse anti-PH3 (1:3000, Abcam 14955), chicken anti-GFP (1:1000, Abcam 13970), rabbit anti-cDCP-1 (1:100, Cell Signaling 9578S), chicken anti-beta-gal (1:1000, Abcam 9361), mouse anti-Achaete (1:100, DSHB). All secondary antibodies used were raised in Donkey, used at 1:500 and purchased by Jackson ImmunoResearch: anti-rat (DyLight™ 405 AffiniPure™ AB_2340681, or Alexa Fluor® 647 AffiniPure™ AB_2340694), anti-mouse (DyLight™ 405 AffiniPure™ AB_2340840, or Alexa Fluor® 488, AffiniPure™ AB_2341099), anti-rabbit (Cy™3 AffiniPure™ AB_2307443, or Alexa Fluor® 647 AffiniPure™ AB_2492288), anti-chicken (Alexa Fluor® 488 AffiniPure™ AB_2340375).

### Pupariation, adult wing dissection, imaging and analysis

Crosses were done by placing 350 virgins and 200 males in egg laying cages with juice-agar plates, either at 18°C (for experiments involving ci-Gal4) or at 25°C (for all other experiments). Plates were changed at 10 am and 6 pm. Freshly hatched L1 larvae were harvested at 6 pm of day *n* from plates changed at 6 pm on day *n-1* (at 25°C), or from plates changed at 6 pm on day *n-2* (at 18°C). 40 L1 larvae were transferred to Drosophila vials with fly food. Larvae of experiments involving ci-Gal4 were kept at 18°C (except for the 8h of apoptosis induction at 29°C). Larvae of all other experiments were kept at 25°C until 48h AEL, when clones were induced by a heat shock (hs) at 37°C. After hs, larvae were kept at 18°C, except for apoptosis induction for 8-16h at 29°C.

#### Pupariation of females only was scored

For adult wing analysis, female flies were fixed in 70% Ethanol. Only the left wing of females was dissected out and mounted dorsal side up on a glass slide in mounting medium (1:1 ethanol 80% and lactic acid 90%). Imaging was performed on a Zeiss Discovery V8 stereomicroscope equipped with a Zeiss Axiocam ICc 5 camera. All wings were imaged using the same parameters and in the same orientation. Wing size analysis was perfomed as described in ^2^, using the following softwares: Fiji, Wings 4 and CPR.

### Image analysis

All 3D stacks were analysed using Fiji. Maximum intensity projection or Local Z projection (Fiji plugin LocalZprojector^57^, E-cadherin as a reference) were performed. PH3-labelled cells were detected using a custom-made Fiji Macro that thresholded the PH3 signal and placed a single pixel dot in the center of each PH3 ROI. Clones were either detected by thresholding the GFP intensity, or manually drawn when thresholding was impossible. The perimeter of the pouch was manually drawn. PH3 index was calculated as follows: PH3 index = (number of PH3 cells/Area of pouch in pixels) x 10 000. Binary images of PH3 dots, clones and pouch were then used for distribution analysis in Python or Matlab.

#### Distribution analysis in Python and Matlab

A custom-made Python routine allowed to automatically detect the border of the clones and draw concentric bands of 5µm around each clone. Bands were set to terminate when reaching the border of the pouch. PH3 density was calculated as the number of PH3 dots divided by band area. This density was normalized to the mean PH3 density (*i.e.* normalized PH3 density) in some Figure panels.

To study whether the observed PH3 distribution differed from a random dot distribution, we randomly distributed dots (same number as the experimental number of PH3 cells) throughout the pouch and computed the associated random dots distribution. Nearest neighbour analysis was performed using Matlab. In brief, for each PH3 dot, the shortest distance to the nearest clone was calculated using the *dsearchn* function.

### Clustering analysis with K and R functions

In order to determine whether the proliferation events exhibit clustering or randomness we are going to apply methods from the spatial statistics for analyzing point patterns. To study clustering, we use Ripley’s functions^24,26^ such as K function, L function, and G function. These functions are designed to analyze the spatial relationships between points and help detect clustering patterns in point processes.

#### K function

Ripley’s K function, is one of the most widely used tools for detecting clustering in point processes. It provides a measure of the average number of points within a given distance *h* of an arbitrary point in the pattern. It compares the observed point pattern to a homogeneous Poisson process to detect clustering or dispersion.

The K function is formally defined as:

Where: λ is the point density of the study region, *n* is the total number of points, and *d_i,j_* is the distance between points *i* and *j*. is an indicator function that equals 1 if the distance is less than or equal to *h*, and 0 otherwise.

For a homogeneous Poisson process points are distributed evenly across the area and the average number of points on a circled area of radius *h* is simply *λπh*^2^, hence the K function values are *K(h)=π h*^2^. When *K(h)>π h*^2^, this indicates some degree of clustering at the scale *h*, and when *K(h)<π h*^2^, this indicates some degree of dispersion at the scale *h*.

#### L function

The L function is a transformation of the K-function that is sometimes easier to interpret. It is defined as:

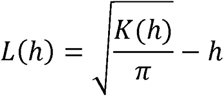

For a homogeneous random process we have *L(h)=*0. When *L(h)>*0 we have clustering at scale *h*.

#### G function

Ripley’s G function keeps track of the proportion of points for which the nearest neighbor is within a given distance threshold, and plots that cumulative percentage against the increasing distance radii, where:

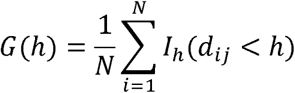

For a Poisson process, G function follows a known theoretical form, and deviations from this form can indicate clustering or dispersion. If the function increases *rapidly* with distance, this is indicative of a clustered pattern. If it increases *slowly* with distance, it is indicated of a dispersed pattern. Something in the middle will be difficult to distinguish from pure chance and is considered random.

#### Wiegand and Moloney (WM) correction

One of the challenges in analyzing point patterns is accounting for edge effects, where points near the boundary of the study region may appear to be less clustered simply because there are fewer neighboring points available due to the boundary constraint. To address this issue we use the corrected K function:

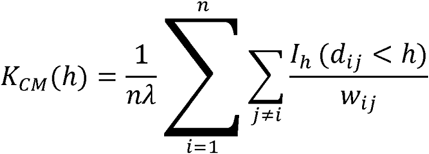

where *w_ij_* is a weight, such that 0 ≤ *w_ij_* ≤ 1 By extension the corrected L function is:

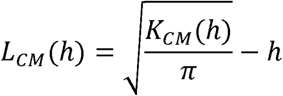

There are many methods that can be used to calculate weights in order to address the edge effect. As we study point patterns on regions (wing pouches) that are never of a regular shape (e.g. a rectangle or a circle), we decided to use Wiegand and Moloney (WM) correction^58^. It is a numerical method, especially adjusted for our case where we work with the real-life images. We work with ‘.tif’ images, where we have stored information on the position of proliferation events, but also on the boundary of the domain of interest, i.e. pouches. For each pixel in the image we have its coordinates and the information weather it belongs to the pouch or outside. The steps to compute the weights for the WM correction are the following:

- Generate a circle, *Ci*(*h*) of a radius *h* with the center at the reference point and store all the coordinates of pixels within the generated circle.
- From the image of the pouch extract a subsection area where we’re sure to find our generated circle.
- Verify which of the pixels of the generated circle *C_i_*(*h*) belong to the pouch, and which remain out of it.
- Finally, calculate the ratio of the number of pixels where the area and the circle overlap, and the total number of pixels in the circle:

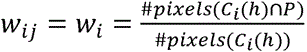, where *P* stands for pouch.

We have used all of the above-mentioned methods in order to detect if there is clustering in our data. If clustering was detected by at least one of the method we would count the wing disc as clustered. If no method detected any clustering, we would classify the distribution as purely random.

We have implemented the clustering analysis in Python, using the standard libraries such as numpy, mathplotlib, scipy and pandas. For manipulation of ‘.tif’ files we used tifffile library. For some simulations of the random Poisson processes and calculation of G functions we used pointpats library (https://pysal.org/pointpats/). For the calculation of K functions, L functions and the boundary correction we developed our own code.

### Statistics

Data were not analysed blindly. No specific method was used to predetermine the number of samples. The definition of n and the number of samples is given in each figure legend and in the table of the Experimental model section. Error bars are standard error of the mean (SEM) or standard deviation (SD). p-values are calculated through t-test if the data passed normality test (Shapiro-Wilk test), Mann-Whitney test/Rank sum test if the distribution was not normal, or one-way ANOVA for multiple comparisons. Statistical tests were performed on Graphpad Prism.

---

**Supplementary Figure 1:**
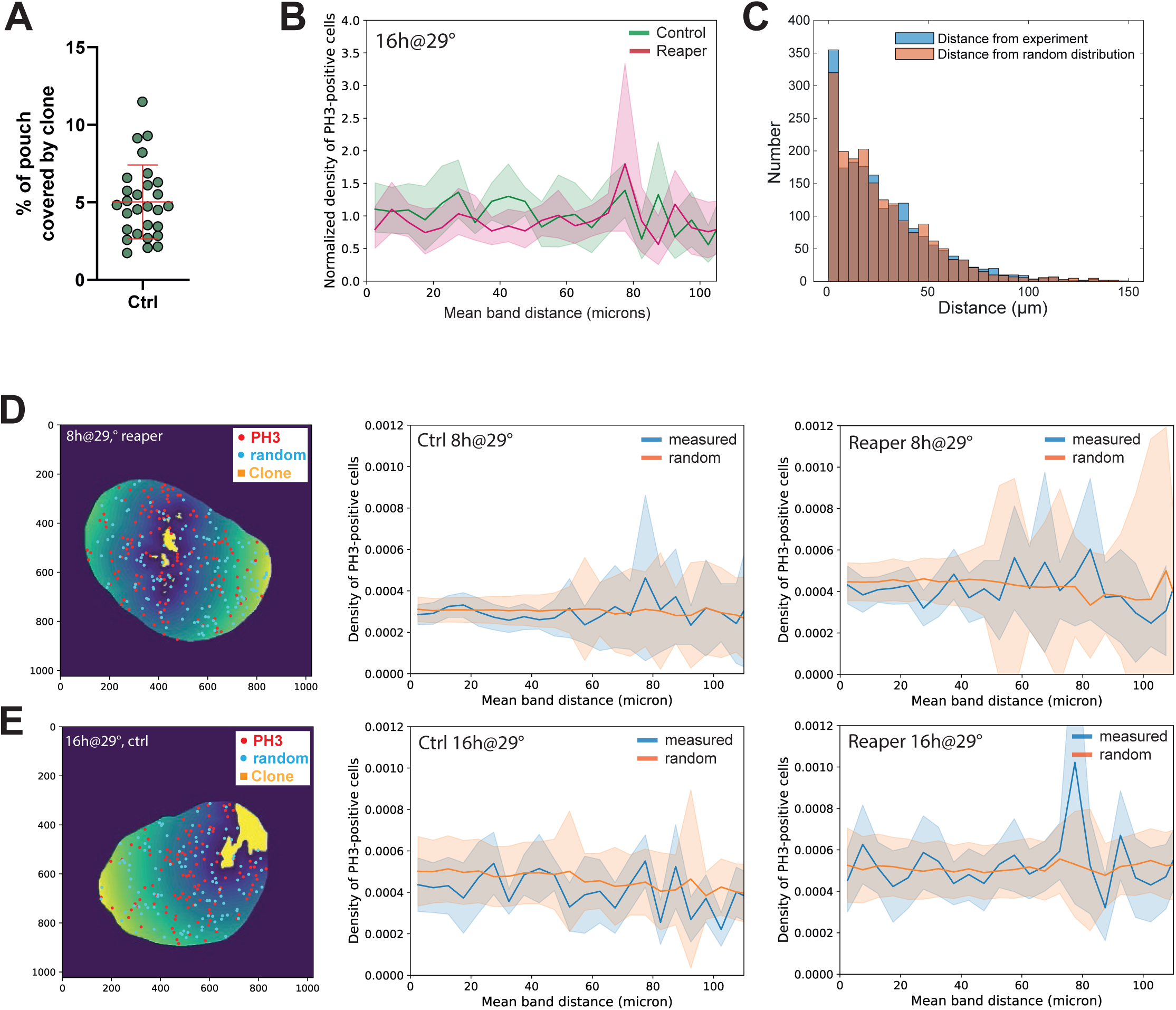
There is no spatial bias in dividing cell distribution near apoptotic clones (associated with Figure 1) **(A)** Percentage of surface coverage of wing pouch by clones. Clones are induced for 10 minutes at L2 stage and allowed to grow at 18°C for 4 days, before UAS-GFP expression for 8h at 29°C in Control condition. n=28 discs from 3 independent experiments. **(B)** Normalised density of PH3-positive cells as a function of the distance of bands, for control (green) or apoptotic (magenta) clones, after 16h of death induction. n=10 discs for Ctrl and n=10 discs for Reaper, from 2 independent experiments. Mean + 95% confidence interval. **(C)** Nearest neighbour analysis. For each proliferating cell, the closest distance to an apoptotic clone is computed, and compared to a randomised distribution of proliferating cells. The histogram shows the distribution of the nearest neighbour distance for experiments with Reaper clones (blue, 16h @29°C) or upon randomisation of the localization of PH3 dividing cells (orange). n=28 discs for Reaper after 8h of apoptosis induction. **(D)** 8h of apoptosis induction. Left: Concentric bands of 5µm (from dark blue to yellow) are drawn around clones (yellow) and the density of proliferating cells (red dots) or randomly distributed cells (cyan) in each band is computed by a custom-made routine in Python. Density of PH3-positive (blue) or random (orange) dots as a function of the mean band distance from Control (middle) or Reaper (right) clones discs. n=28 discs for Ctrl and n=18 discs for Reaper, from 3 independent experiments. Mean +/−0.5 SD **(E)** 16h of apoptosis induction. Left: Concentric bands of 5µm (from dark blue to yellow) are drawn around clones (yellow) and the density of proliferating cells (red dots) or randomly distributed cells (cyan) in each band is computed by a custom-made routine in Python. Density of PH3-positive (blue) or random (orange) dots as a function of the mean band distance for Control (middle) or Reaper (right) clones discs. n=10 discs for Ctrl and n=10 discs for Reaper, from 2 independent experiments. Mean +/− 0.5 SD

**Supplementary Figure 2:**
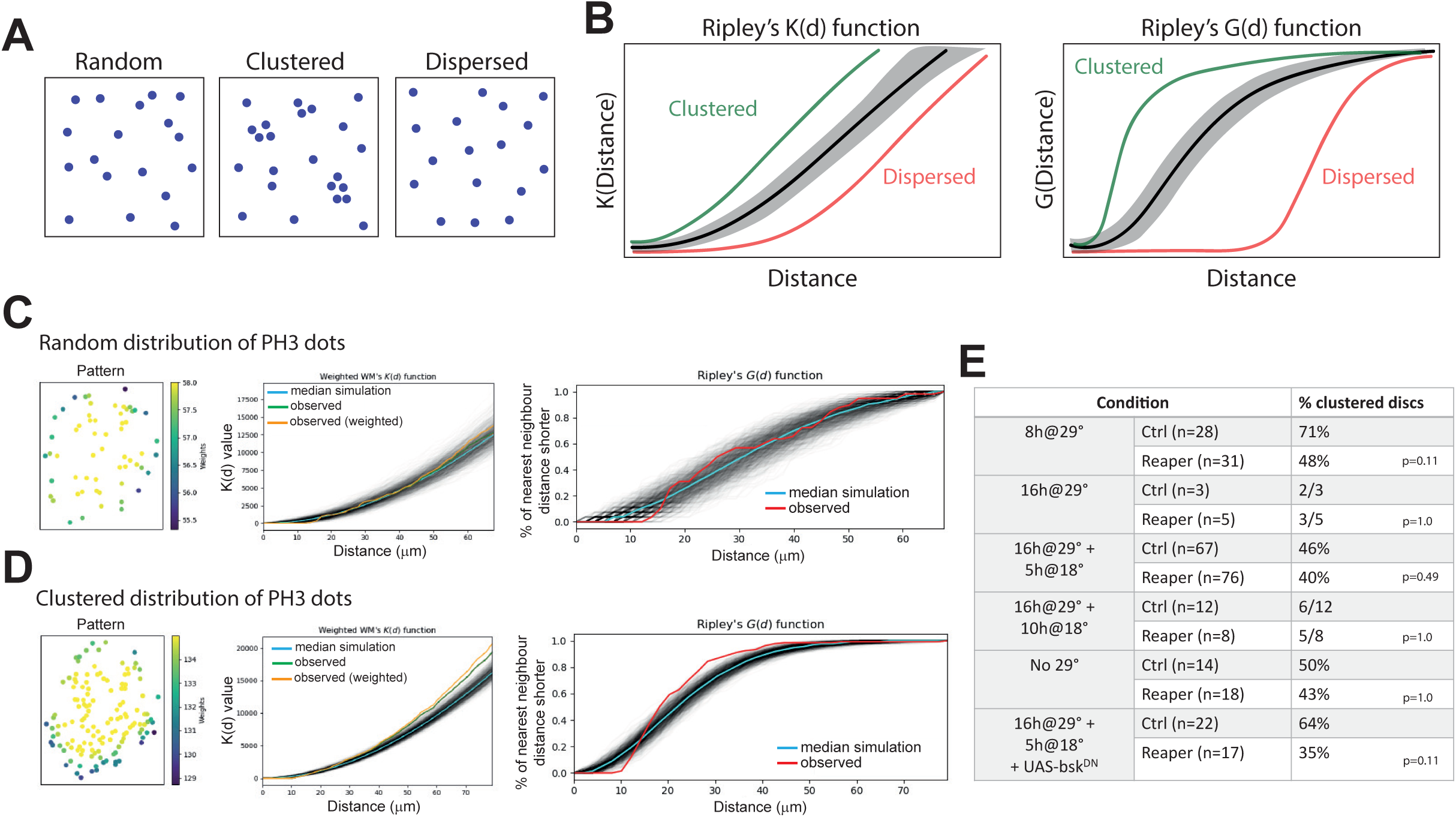
Ripley function analysis of PH3 cells spatial distribution (associated with Figure 1) **(A)** Examples of random (left), clustered (middle) and dispersed (right) point distributions. **(B)** Schematics of Ripley’s K (left) and G (right) functions. Gray: median of the simulation of a Poisson process (random), red: curve for a dispersed point distribution, green: curve for a clustered point distribution. **(C)** Example of random distribution of PH3-stained cells from one wing disc. Left: Distribution of PH3-stained cells in the pouch, with weights associated with the proximity of cells to the border of the pouch, used to correct biases related to boundary effects (see **Methods**). Middle: K-function, right: G-function, grey curves are single simulation of distribution using a Poisson process, blue curve the median of the simulation, green curve the raw experimental distribution, orange the experimental distribution corrected with the weights. **(D)** Example of clustered distribution of PH3-stained cells from one wing disc. Left: Distribution of PH3-stained cells in the pouch, with weights associated with the proximity of cells to the border of the pouch, used to correct biases related to boundary effects (see **Methods**). Middle: K-function, right: G-function, grey curves are single simulation of distribution using a Poisson process, blue curve the median of the simulation, green curve the raw experimental distribution, orange the experimental distribution corrected with the weights. **(E)** Table summarising the percentages of discs with a clustered distributions of PH3 cells throughout the different experimental conditions using the K-function and G-function (n, number of discs). PH3 distribution was counted as clustered if it showed significant clustering in at least one of the methodology (K or G function, see **Methods**). Note that we noticed a tendency to have more clustered distribution at later developmental stage where a pattern of PH3 appear near D/V boundary. p values compare the proportion of clustered discs between control and Reaper with Fisher’s exact test (no significant differences).

**Supplementary Figure 3:**
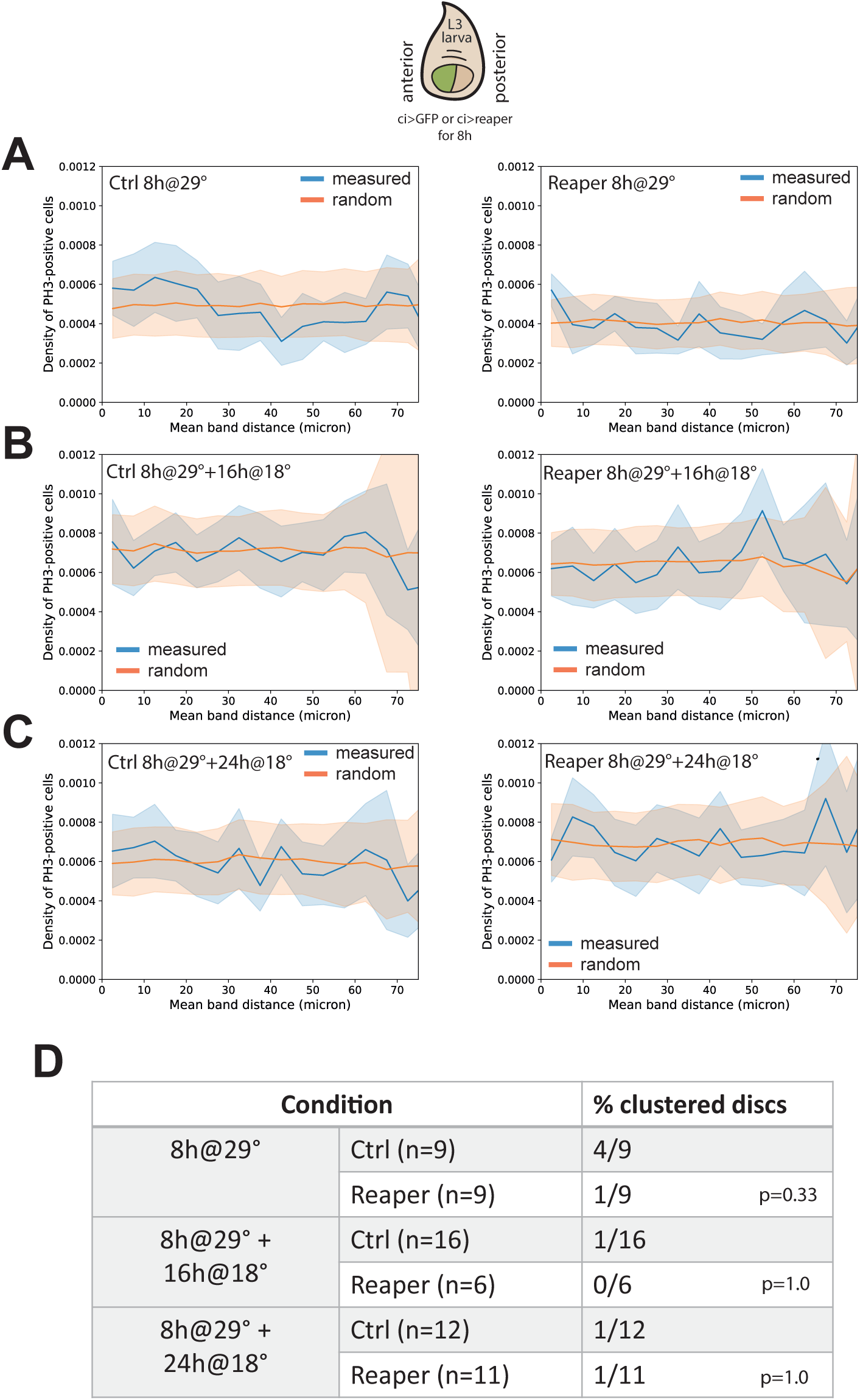
Analysis of mitotic cell distribution in the posterior compartment upon induction of apoptosis in the anterior compartment (associated with Figure 1) **(A)** Density of PH3-positive (blue) or random (orange) dots in the posterior domain as a function of the band distance to the anterior compartment for Control (left) or Reaper (right) discs, for 8h of apoptosis induction in the anterior (ci) domain. n=17 discs for Ctrl and n=17 discs for Reaper, from 2 independent experiments. Mean +/−0.5 SD **(B)** Density of PH3-positive (blue) or random (orange) dots in the posterior domain as a function of the band distance to the anterior compartment for Control (left) or Reaper (right) discs, for 8h of apoptosis induction in the anterior (ci) domain followed by 16h or recovery. n=24 discs for Ctrl and n=14 discs for Reaper, from 2 independent experiments. Mean +/−0.5 SD **(C)** Density of PH3-positive (blue) or random (orange) dots in the posterior domain as a function of the band distance to the anterior compartment for Control (left) or Reaper (right) discs, for 8h of apoptosis induction in the anterior (ci) domain followed by 24h or recovery. n=20 discs for Ctrl and n=19 discs for Reaper, from 2 independent experiments. Mean +/−0.5 SD **(D)** Table summarising the percentages of discs (control or upon apoptosis induction in the ci domain) with a clustered distribution of PH3 cells using the K-function and G-function (see **Fig. S2, Methods**). p values compare the proportion of clustered discs between control and Reaper with Fisher’s exact test (no significant difference).

**Supplementary Figure 4:**
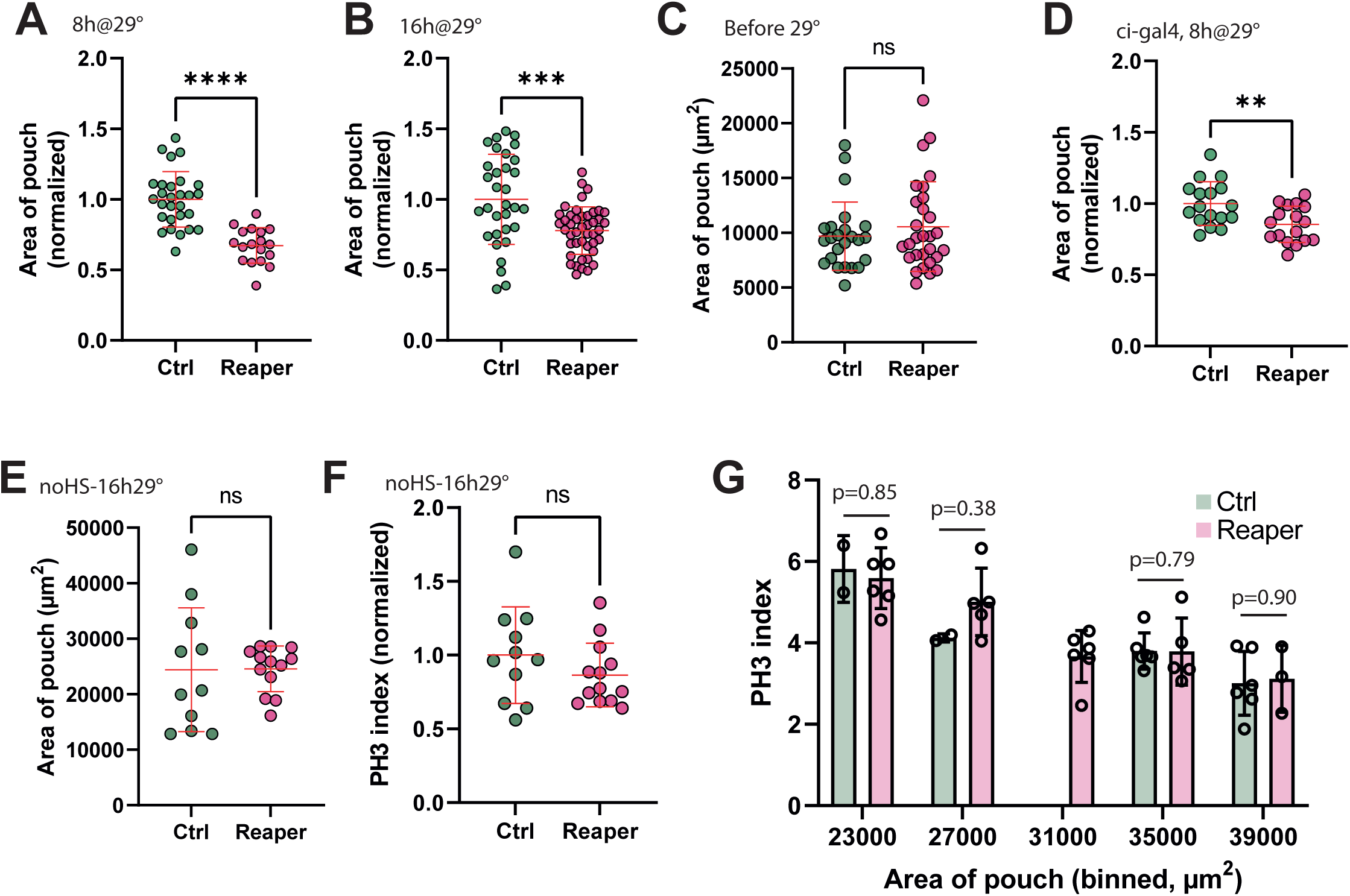
Apoptosis induction in clone reduces transiently disc size (associated with Figure 2) **(A)** Area of the pouch (normalised) after 8h of apoptosis induction in clones for Ctrl (n=28, green, 3 independent experiments) and Reaper (n=18, magenta, 3 independent experiments). Welch’s t test, p<0.0001 ****. **(B)** Area of the pouch (normalised) after 16h of apoptosis induction in clones for Ctrl (n=32, green, 2 independent experiments) and Reaper (n=44, magenta, 2 independent experiments). Welch’s t test, p<0.001 ***. **(C)** Area of the pouch at 96h AEL, before apoptosis induction for Ctrl (n=24, green, 2 independent experiments) and Reaper (n=29, magenta, 2 independent experiments), without any incubation at the permissive temperature. Mann-Whitney test, p=0.6259. **(D)** Area of the pouch (normalised) after 8h of apoptosis induction in the anterior (ci) domain for Ctrl (n=17, green, 2 independent experiments) and Reaper (n=17, magenta, 2 independent experiments). Welch’s t test, p<0.01 **. **(E)** Larvae are transferred to 29°C at 96h AEL, and dissected after 16h at 29°C. The graph shows the area of the pouch in the absence of clone induction (no heat shock) for Ctrl (n=11, green, 1 experiment) and Reaper (n=13, magenta, 1 experiment) to check for potential interaction between temperature and a putative genetic background effect. Mann-Whitney test, p=0.6905. **(F)** Normalised PH3 index after 16h at 29°C (for the same individuals as **Fig. S4, E**), in the absence of clone induction (no heat shock) for Ctrl (n=11, green, 1 experiment) and Reaper (n=13, magenta, 1 experiment). Mann-Whitney test, p=0.3918. **(G)** PH3 index as a function of binned values of the pouch area for Control (green) and Reaper (magenta), after either 8h or 16h of apoptosis induction. n=16 discs for Ctrl (5 independent experiments) and n=25 discs for Reaper (5 independent experiments). When discs of similar size are compared, there is no difference in the PH3 index. P values are Mann-Whitney tests.

**Supplementary Figure 5:**
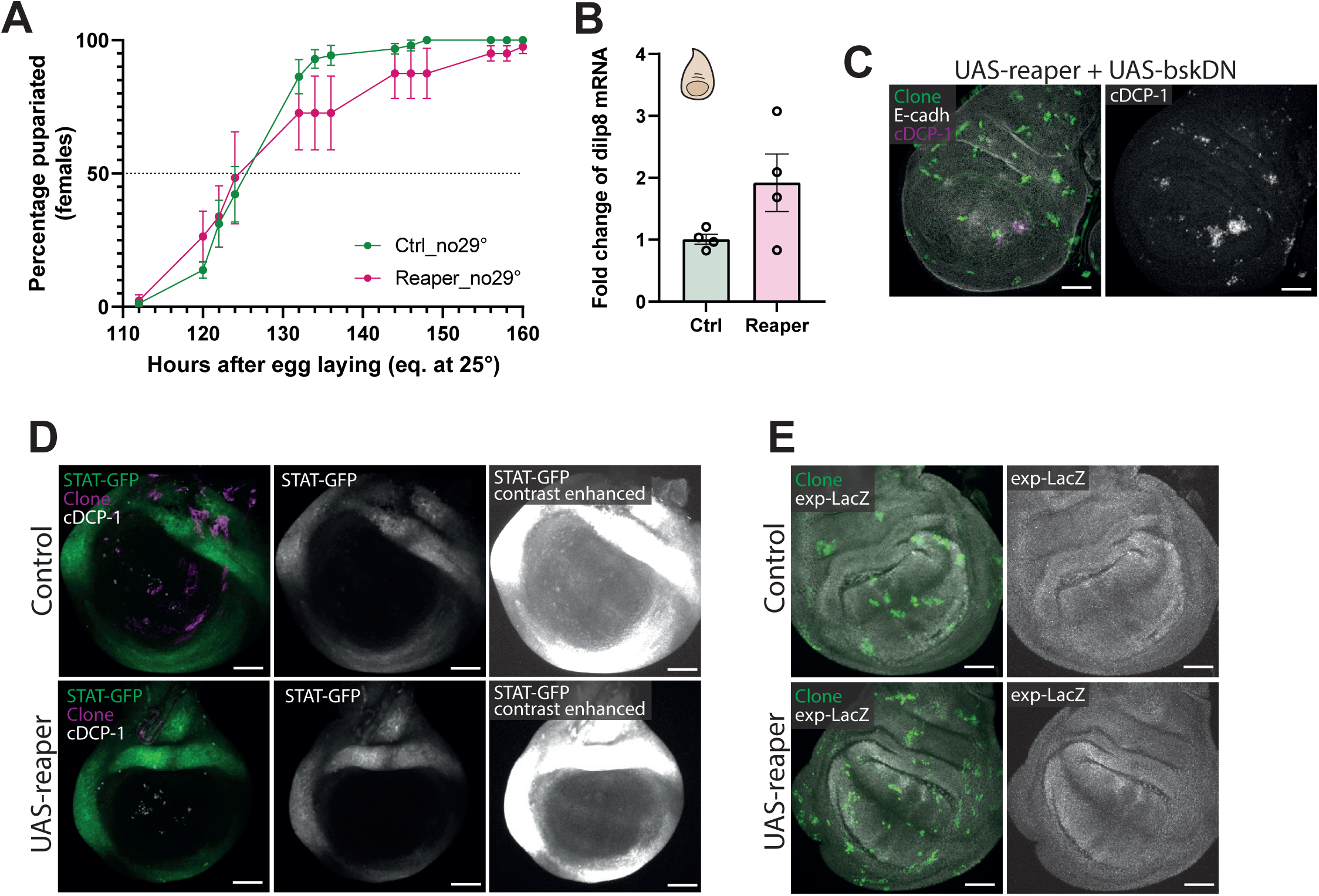
JNK activation in clones is responsible for pupariation delay (associated with. **Figure 3). (A)** Percentage of pupariated females (only females have both UAS-reaper and hs-flp) for Control (green) and Reaper (magenta) individuals. Clones are induced by a heath shock at 48h AEL, but not allowed to undergo apoptosis (tubes are kept at 18°C after heath shock). n=68 females (Control) and n= 41 females (Reaper) from 2 independent experiments. Mean+/− SEM. **(B)** Fold change of dilp8 mRNA levels by qPCR from whole wing discs. Clones are induced at 48h AEL, allowed to grow at 18°C until 96h AEL and then switch to 29°C for 16h to induce apoptosis (or GFP for controls), after which discs are dissected. n= approximatively 60 discs for Ctrl and n= approximatively 60 discs for reaper, from 4 independent experiments (one dot per experiment). **(C)** Inhibition of JNK does not suppress apoptosis in clones. Clones (green, act<cd2<Gal4, UAS-GFP x UAS-reaper, UAS-bskDN), E-cadh (white), cDCP-1 (cleaved caspase, magenta and white). Scale bars, 50µm. **(D)** Apoptosis induction in clones does not lead to STAT activation. Clones (magenta), cDCP-1 (white), STAT-GFP (green and white, enhanced contrast on the right). Scale bars, 50µm. **(E)** Apoptosis induction in clones does not lead to expanded activation (expanded is a readout of Hippo pathway). Clones (green), expanded-LacZ (white). Scale bars, 50µm.

**Supplementary Figure 6:**
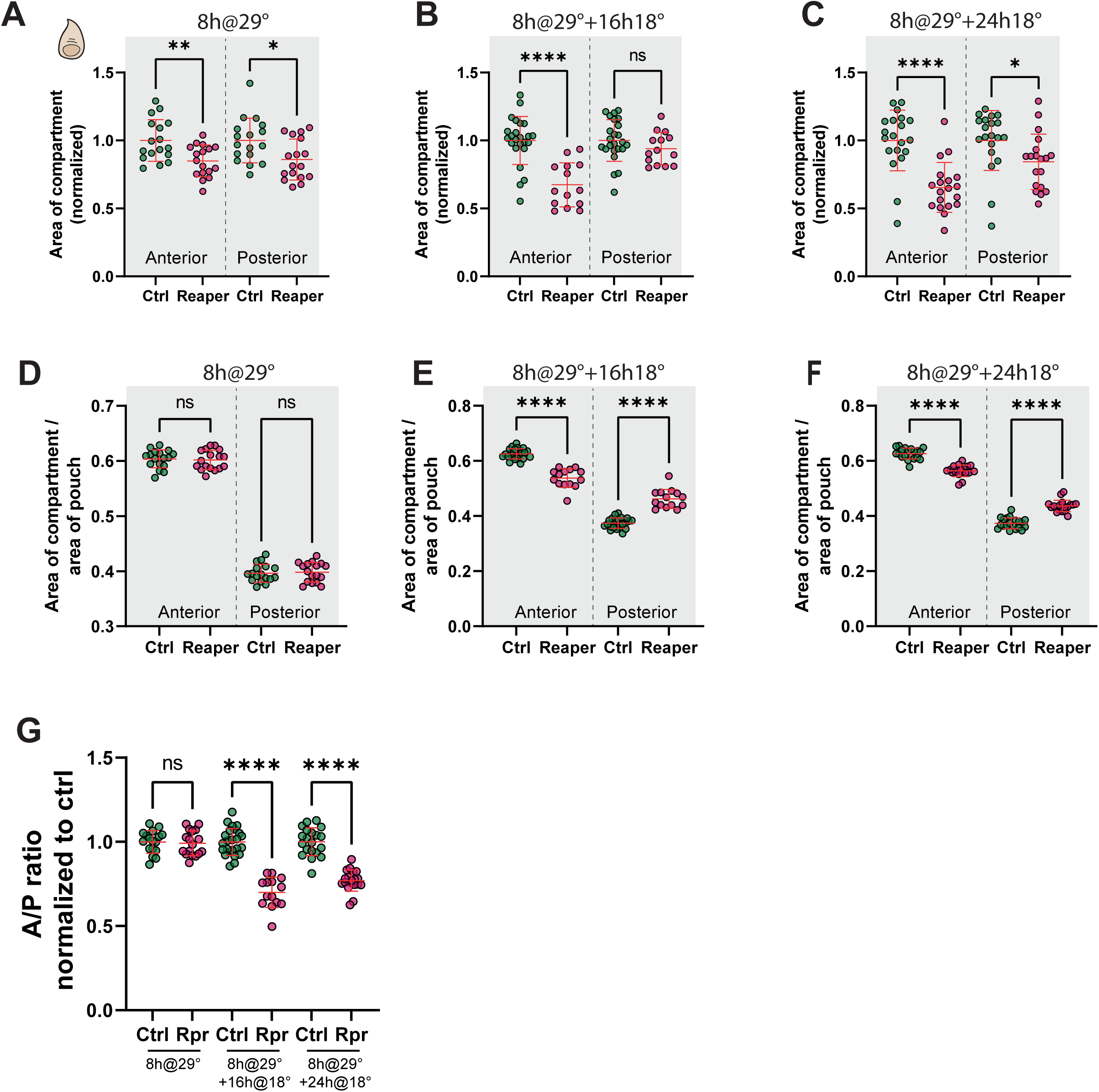
Induction of apoptosis in the anterior compartment reduces the relative size of the compartment throughout larval stage. **(A)** Area of the anterior (A) or posterior (P) compartment in the larva, normalized to the Ctrl corresponding compartment, after 8h of apoptosis induction in the Ci domain (anterior) at 92h AEL. Ctrl (n=17 discs, 2 independent experiments), Reaper (n=17 discs, 2 independent experiments. Mean+/−SD, one-way ANOVA, p<0.05 *, p<0.01 **. **(B)** Area of the anterior (A) or posterior (P) compartment in the larva, normalized to the Ctrl corresponding compartment, after 8h of apoptosis induction in the Ci domain (anterior) at 80h AEL followed by 16h of recovery. Ctrl (n=24 discs, 2 independent experiments), Reaper (n=14 discs, 2 independent experiments. Mean+/−SD, one-way ANOVA, p<0.0001 ****. **(C)** Area of the anterior (A) or posterior (P) compartment in the larva, normalized to the Ctrl corresponding compartment, after 8h of apoptosis induction in the Ci domain (anterior) at 80h AEL followed by 24h of recovery. Ctrl (n=20 discs, 2 independent experiments), Reaper (n=19 discs, 2 independent experiments. Mean+/−SD, one-way ANOVA, p<0.0001 ****. **(D)** Area of A or P compartment divided by the area of the pouch after 8h of apoptosis induction in the Ci domain (anterior) at 92h AEL. Ctrl (n=17 discs, 2 independent experiments), Reaper (n=17 discs, 2 independent experiments. Mean+/−SD, one-way ANOVA, ns, not significant. **(E)** Area of A or P compartment divided by the area of the pouch after 8h of apoptosis induction in the Ci domain (anterior) at 80h AEL followed by 16h of recovery. Ctrl (n=24 discs, 2 independent experiments), Reaper (n=14 discs, 2 independent experiments. Mean+/−SD, one-way ANOVA, p<0.0001 ****. **(F)** Area of A or P compartment divided by the area of the pouch after 8h of apoptosis induction in the Ci domain (anterior) at 80h AEL followed by 24h of recovery. Ctrl (n=20 discs, 2 independent experiments), Reaper (n=19 discs, 2 independent experiments. Mean+/−SD, one-way ANOVA, p<0.0001 ****. **(G)** A/P ratio of disc upon induction of apoptosis in the Ci (anterior) compartment for 8h, normalized to the A/P ratio of Ctrl at the given time-points, from the same discs as the one used in **Fig. S6, A-F**. Mean+/−SD, one-way ANOVA, p<0.0001 ****.

## References

1. Watson, A.J.M., Duckworth, C.A., Guan, Y.F., and Montrose, M.H. (2009). Mechanisms of Epithelial Cell Shedding in the Mammalian Intestine and Maintenance of Barrier Function. Molecular Structure and Function of the Tight Junction: From Basic Mechanisms to Clinical Manifestations 1165, 135–142. 10.1111/j.1749-6632.2009.04027.x.

2. Matamoro-Vidal, A., Cumming, T., Davidovic, A., Levillayer, F., and Levayer, R. (2024). Patterned apoptosis has an instructive role for local growth and tissue shape regulation in a fast-growing epithelium. Curr Biol 34, 376–388 e377. 10.1016/j.cub.2023.12.031.

3. Boulan, L., and Léopold, P. (2021). What determines organ size during development and regeneration? Development (Cambridge, England) 148. 10.1242/dev.196063.

4. Boulan, L., Milán, M., and Léopold, P. (2015). The systemic control of growth. Cold Spring Harbor Perspectives in Biology 7. 10.1101/cshperspect.a019117.

5. Hariharan, I.K. (2015). Organ Size Control: Lessons from Drosophila. Dev Cell 34, 255–265. 10.1016/j.devcel.2015.07.012.

6. Fan, Y., and Bergmann, A. (2008). Apoptosis-induced compensatory proliferation. The Cell is dead. Long live the Cell! Trends Cell Biol 18, 467–473. 10.1016/j.tcb.2008.08.001.

7. Martin, F.A., Perez-Garijo, A., and Morata, G. (2009). Apoptosis in Drosophila: compensatory proliferation and undead cells. Int J Dev Biol 53, 1341–1347. 10.1387/ijdb.072447fm.

8. Haynie, J.L., and Bryant, P.J. (1977). The effects of X-rays on the proliferation dynamics of cells in the imaginal wing disc ofDrosophila melanogaster. Wilehm Roux Arch Dev Biol 183, 85–100. 10.1007/BF00848779.

9. Verghese, S., and Su, T.T. (2017). STAT, Wingless, and Nurf-38 determine the accuracy of regeneration after radiation damage in Drosophila. PLoS Genet 13, e1007055. 10.1371/journal.pgen.1007055.

10. Smith-Bolton, R.K., Worley, M.I., Kanda, H., and Hariharan, I.K. (2009). Regenerative growth in Drosophila imaginal discs is regulated by Wingless and Myc. Dev Cell 16, 797–809. 10.1016/j.devcel.2009.04.015.

11. Ursprung, H., and Hadorn, E. (1962). [Further research on model growth in combination with partly dissociated wing imaginal disks of Drosophila melanogaster]. Dev Biol 4, 40–66. 10.1016/0012-1606(62)90032-5.

12. Diaz-Garcia, S., and Baonza, A. (2013). Pattern reorganization occurs independently of cell division during Drosophila wing disc regeneration in situ. Proc Natl Acad Sci U S A 110, 13032–13037. 10.1073/pnas.1220543110.

13. Perez-Garijo, A., Martin, F.A., Struhl, G., and Morata, G. (2005). Dpp signaling and the induction of neoplastic tumors by caspase-inhibited apoptotic cells in Drosophila. Proc Natl Acad Sci U S A 102, 17664–17669. 10.1073/pnas.0508966102.

14. Fan, Y., and Bergmann, A. (2008). Distinct mechanisms of apoptosis-induced compensatory proliferation in proliferating and differentiating tissues in the Drosophila eye. Dev Cell 14, 399–410. 10.1016/j.devcel.2008.01.003.

15. Perez-Garijo, A., Martin, F.A., and Morata, G. (2004). Caspase inhibition during apoptosis causes abnormal signalling and developmental aberrations in Drosophila. Development 131, 5591–5598. 10.1242/dev.01432.

16. Cosolo, A., Jaiswal, J., Csordas, G., Grass, I., Uhlirova, M., and Classen, A.K. (2019). JNK-dependent cell cycle stalling in G2 promotes survival and senescence-like phenotypes in tissue stress. Elife 8. 10.7554/eLife.41036.

17. Jaiswal, J., Egert, J., Engesser, R., Peyroton, A.A., Nogay, L., Weichselberger, V., Crucianelli, C., Grass, I., Kreutz, C., Timmer, J., and Classen, A.K. (2023). Mutual repression between JNK/AP-1 and JAK/STAT stratifies senescent and proliferative cell behaviors during tissue regeneration. PLoS Biol 21, e3001665. 10.1371/journal.pbio.3001665.

18. Chera, S., Ghila, L., Dobretz, K., Wenger, Y., Bauer, C., Buzgariu, W., Martinou, J.C., and Galliot, B. (2009). Apoptotic cells provide an unexpected source of Wnt3 signaling to drive hydra head regeneration. Dev Cell 17, 279–289. 10.1016/j.devcel.2009.07.014.

19. Jiang, H., Grenley, M.O., Bravo, M.J., Blumhagen, R.Z., and Edgar, B.A. (2011). EGFR/Ras/MAPK signaling mediates adult midgut epithelial homeostasis and regeneration in Drosophila. Cell Stem Cell 8, 84–95. 10.1016/j.stem.2010.11.026.

20. Tseng, A.S., Adams, D.S., Qiu, D., Koustubhan, P., and Levin, M. (2007). Apoptosis is required during early stages of tail regeneration in Xenopus laevis. Dev Biol 301, 62–69. 10.1016/j.ydbio.2006.10.048.

21. Brock, C.K., Wallin, S.T., Ruiz, O.E., Samms, K.M., Mandal, A., Sumner, E.A., and Eisenhoffer, G.T. (2019). Stem cell proliferation is induced by apoptotic bodies from dying cells during epithelial tissue maintenance. Nature Communications 2019 10:1 10, 1–11. 10.1038/s41467-019-09010-6.

22. Ankawa, R., Goldberger, N., Yosefzon, Y., Koren, E., Yusupova, M., Rosner, D., Feldman, A., Baror-Sebban, S., Buganim, Y., Simon, D.J., et al. (2021). Apoptotic cells represent a dynamic stem cell niche governing proliferation and tissue regeneration. Dev Cell 56, 1900–1916 e1905. 10.1016/j.devcel.2021.06.008.

23. Kawaue, T., Yow, I., Pan, Y., Le, A.P., Lou, Y., Loberas, M., Shagirov, M., Teng, X., Prost, J., Hiraiwa, T., et al. (2023). Inhomogeneous mechanotransduction defines the spatial pattern of apoptosis-induced compensatory proliferation. Dev Cell 58, 267–277 e265. 10.1016/j.devcel.2023.01.005.

24. Ripley, B.D. (1976). The second-order analysis of stationary point processes. Journal of Applied Probability 13, 255–266. 10.2307/3212829.

25. Valon, L., Davidovic, A., Levillayer, F., Villars, A., Chouly, M., Cerqueira-Campos, F., and Levayer, R. (2021). Robustness of epithelial sealing is an emerging property of local ERK feedback driven by cell elimination. Dev Cell 56, 1700–1711 e1708. 10.1016/j.devcel.2021.05.006.

26. Ripley, B.D. (1988). Statistical inference for spatial processes (Cambridge University Press).

27. Cubas, P., de Celis, J.F., Campuzano, S., and Modolell, J. (1991). Proneural clusters of achaete-scute expression and the generation of sensory organs in the Drosophila imaginal wing disc. Genes Dev 5, 996–1008. 10.1101/gad.5.6.996.

28. Nogueira Alves, A., Oliveira, M.M., Koyama, T., Shingleton, A., and Mirth, C.K. (2022). Ecdysone coordinates plastic growth with robust pattern in the developing wing. Elife 11. 10.7554/eLife.72666.

29. Martin, F.A., Herrera, S.C., and Morata, G. (2009). Cell competition, growth and size control in the Drosophila wing imaginal disc. Development 136, 3747–3756. 10.1242/dev.038406.

30. Strassburger, K., Lutz, M., Muller, S., and Teleman, A.A. (2021). Ecdysone regulates Drosophila wing disc size via a TORC1 dependent mechanism. Nat Commun 12, 6684. 10.1038/s41467-021-26780-0.

31. Stieper, B.C., Kupershtok, M., Driscoll, M.V., and Shingleton, A.W. (2008). Imaginal discs regulate developmental timing in Drosophila melanogaster. Dev Biol 321, 18–26. 10.1016/j.ydbio.2008.05.556.

32. Colombani, J., Andersen, D.S., and Leopold, P. (2012). Secreted peptide Dilp8 coordinates Drosophila tissue growth with developmental timing. Science 336, 582–585. 10.1126/science.1216689.

33. Ryoo, H.D., Gorenc, T., and Steller, H. (2004). Apoptotic cells can induce compensatory cell proliferation through the JNK and the Wingless signaling pathways. Dev Cell 7, 491–501. 10.1016/j.devcel.2004.08.019.

34. Fan, Y., Wang, S., Hernandez, J., Yenigun, V.B., Hertlein, G., Fogarty, C.E., Lindblad, J.L., and Bergmann, A. (2014). Genetic models of apoptosis-induced proliferation decipher activation of JNK and identify a requirement of EGFR signaling for tissue regenerative responses in Drosophila. PLoS Genet 10, e1004131. 10.1371/journal.pgen.1004131.

35. Perez-Garijo, A., Shlevkov, E., and Morata, G. (2009). The role of Dpp and Wg in compensatory proliferation and in the formation of hyperplastic overgrowths caused by apoptotic cells in the Drosophila wing disc. Development 136, 1169–1177. 10.1242/dev.034017.

36. Bergantinos, C., Corominas, M., and Serras, F. (2010). Cell death-induced regeneration in wing imaginal discs requires JNK signalling. Development 137, 1169–1179. 10.1242/dev.045559.

37. Garelli, A., Gontijo, A.M., Miguela, V., Caparros, E., and Dominguez, M. (2012). Imaginal discs secrete insulin-like peptide 8 to mediate plasticity of growth and maturation. Science 336, 579–582. 10.1126/science.1216735.

38. Colombani, J., Andersen, D.S., Boulan, L., Boone, E., Romero, N., Virolle, V., Texada, M., and Leopold, P. (2015). Drosophila Lgr3 Couples Organ Growth with Maturation and Ensures Developmental Stability. Curr Biol 25, 2723–2729. 10.1016/j.cub.2015.09.020.

39. Garelli, A., Heredia, F., Casimiro, A.P., Macedo, A., Nunes, C., Garcez, M., Dias, A.R.M., Volonte, Y.A., Uhlmann, T., Caparros, E., et al. (2015). Dilp8 requires the neuronal relaxin receptor Lgr3 to couple growth to developmental timing. Nat Commun 6, 8732. 10.1038/ncomms9732.

40. Vallejo, D.M., Juarez-Carreno, S., Bolivar, J., Morante, J., and Dominguez, M. (2015). A brain circuit that synchronizes growth and maturation revealed through Dilp8 binding to Lgr3. Science 350, aac6767. 10.1126/science.aac6767.

41. Chatterjee, N., and Bohmann, D. (2012). A versatile PhiC31 based reporter system for measuring AP-1 and Nrf2 signaling in Drosophila and in tissue culture. PLoS One 7, e34063. 10.1371/journal.pone.0034063.

42. Pinal, N., Ruiz-Losada, M., Azpiazu, N., and Morata, G. (2023). Size compensation in Drosophila after generalised cell death. Front Cell Dev Biol 11, 1301913. 10.3389/fcell.2023.1301913.

43. Liang, J., Balachandra, S., Ngo, S., and O’Brien, L.E. (2017). Feedback regulation of steady-state epithelial turnover and organ size. Nature 548, 588–591. 10.1038/nature23678.

44. Amcheslavsky, A., Lindblad, J.L., and Bergmann, A. (2020). Transiently “Undead” Enterocytes Mediate Homeostatic Tissue Turnover in the Adult Drosophila Midgut. Cell Rep 33, 108408. 10.1016/j.celrep.2020.108408.

45. Martin, J.L., Sanders, E.N., Moreno-Roman, P., Jaramillo Koyama, L.A., Balachandra, S., Du, X., and O’Brien, L.E. (2018). Long-term live imaging of the Drosophila adult midgut reveals real-time dynamics of division, differentiation and loss. Elife 7. 10.7554/eLife.36248.

46. Levayer, R. (2024). Staying away from the breaking point: Probing the limits of epithelial cell elimination. Curr Opin Cell Biol 86, 102316. 10.1016/j.ceb.2023.102316.

47. de Vreede, G., Gerlach, S.U., and Bilder, D. (2022). Epithelial monitoring through ligand-receptor segregation ensures malignant cell elimination. Science 376, 297–301. 10.1126/science.abl4213.

48. Dye, N.A., Popovic, M., Spannl, S., Etournay, R., Kainmuller, D., Ghosh, S., Myers, E.W., Julicher, F., and Eaton, S. (2017). Cell dynamics underlying oriented growth of the Drosophila wing imaginal disc. Development 144, 4406–4421. 10.1242/dev.155069.

49. Herboso, L., Oliveira, M.M., Talamillo, A., Perez, C., Gonzalez, M., Martin, D., Sutherland, J.D., Shingleton, A.W., Mirth, C.K., and Barrio, R. (2015). Ecdysone promotes growth of imaginal discs through the regulation of Thor in D. melanogaster. Sci Rep 5, 12383. 10.1038/srep12383.

50. Perez-Mockus, G., Cocconi, L., Alexandre, C., Aerne, B., Salbreux, G., and Vincent, J.P. (2023). The Drosophila ecdysone receptor promotes or suppresses proliferation according to ligand level. Dev Cell 58, 2128–2139 e2124. 10.1016/j.devcel.2023.08.032.

51. Sanchez, J.A., Mesquita, D., Ingaramo, M.C., Ariel, F., Milan, M., and Dekanty, A. (2019). Eiger/TNFalpha-mediated Dilp8 and ROS production coordinate intra-organ growth in Drosophila. PLoS Genet 15, e1008133. 10.1371/journal.pgen.1008133.

52. Mesquita, D., Dekanty, A., and Milan, M. (2010). A dp53-dependent mechanism involved in coordinating tissue growth in Drosophila. PLoS Biol 8, e1000566. 10.1371/journal.pbio.1000566.

53. Baker, N.E. (2017). Mechanisms of cell competition emerging from Drosophila studies. Curr Opin Cell Biol 48, 40–46. 10.1016/j.ceb.2017.05.002.

54. Matamoro-Vidal, A., and Levayer, R. (2019). Multiple Influences of Mechanical Forces on Cell Competition. Curr Biol 29, R762–R774. 10.1016/j.cub.2019.06.030.

55. Wagstaff, L., Kolahgar, G., and Piddini, E. (2013). Competitive cell interactions in cancer: a cellular tug of war. Trends Cell Biol 23, 160–167. 10.1016/j.tcb.2012.11.002.

56. de la Cova, C., Abril, M., Bellosta, P., Gallant, P., and Johnston, L.A. (2004). Drosophila myc regulates organ size by inducing cell competition. Cell 117, 107–116. S0092867404002144 [pii].

57. Herbert, S., Valon, L., Mancini, L., Dray, N., Caldarelli, P., Gros, J., Esposito, E., Shorte, S.L., Bally-Cuif, L., Aulner, N., et al. (2021). LocalZProjector and DeProj: a toolbox for local 2D projection and accurate morphometrics of large 3D microscopy images. BMC Biol 19, 136. 10.1186/s12915-021-01037-w.

58. Wiegand, T., and A. Moloney, K. (2004). Rings, circles, and null-models for point pattern analysis in ecology. Oikos 104, 209–229. 10.1111/j.0030-1299.2004.12497.x.

